# A schizophrenia risk gene, *NRGN*, bidirectionally modulates synaptic plasticity via regulating the neuronal phosphoproteome

**DOI:** 10.1101/481291

**Authors:** Hongik Hwang, Matthew J. Szucs, Lei J. Ding, Andrew Allen, Henny Haensgen, Fan Gao, Arturo Andrade, Jennifer Q. Pan, Steven A. Carr, Rushdy Ahmad, Weifeng Xu

## Abstract

*NRGN* is a schizophrenia risk gene identified in recent genetic studies, encoding a small neuronal protein, neurogranin (Ng). Individuals carrying a risk variant of *NRGN* showed decreased hippocampal activation during contextual fear conditioning. Furthermore, the expression of Ng was reduced in the post-mortem brains of schizophrenic patients. Using the mouse model, we found that the translation of Ng in hippocampus is rapidly increased in response to novel context exposure, and this up-regulation is required for encoding contextual memory. The extent and degree of the effect that altered Ng expression has on neuronal cellular functions are largely unknown. Here, we found that Ng bidirectionally regulates synaptic plasticity in the hippocampus. Elevated Ng levels facilitated long-term potentiation (LTP), whereas decreased Ng levels impaired LTP. Quantitative phosphoproteomic analysis revealed that decreasing Ng caused a significant shift in the phosphorylation status of postsynaptic density proteins, highlighting clusters of schizophrenia- and autism-related genes. In particular, decreasing Ng led to the hypo-phosphorylation of NMDAR subunit Grin2A at newly identified sites, resulting in accelerated decay of NMDAR-mediated channel currents. blocking protein phosphatase PP2B activity rescued the accelerated synaptic NMDAR current decay and the impairment of LTP caused by decreased Ng levels, suggesting that enhanced synaptic PP2B activity led to the deficits. Taken together, our work suggests that altered Ng levels under pathological conditions affect the phosphorylation status of neuronal proteins by tuning PP2B activity and thus the induction of synaptic plasticity, revealing a novel mechanistic link of a schizophrenia risk gene to cognitive deficits.

## Introduction

Schizophrenia, affecting about 1% of the population worldwide, is a chronic mental disorder with psychotic symptoms, such as delusions, hallucinations and disorganized thinking^1^. Schizophrenia is one of the top leading causes of disability worldwide^2^, and affected individuals suffer from difficulty in social relationships, motor impairment, and cognitive dysfunction, which severely interferes with the patients’ daily functioning. Individuals with schizophrenia have an increased risk of premature mortality, and about 5% of the patients die by suicide^3^. Antipsychotic medications are commonly used to ease psychotic symptoms. However, available symptomatic treatments are only partially effective, and lifelong treatment is required.

A combination of physical, genetic, psychological and environmental factors is thought to contribute to the development of schizophrenia, but the exact pathophysiology is unknown. However, the highly heritable nature of schizophrenia implies a significant role of inherited genetic variants in the etiology^4, 5^. Genome-wide association studies reported more than 100 genetic loci associated with schizophrenia^6-14^, and the neurogranin (Ng, gene name: *NRGN*) gene has been identified as one of schizophrenia risk genes with top associations in different patients population across the world^6, 7, 15-17^. rs12807809 is a single-nucleotide polymorphism located 3,457 bases upstream from the promoter region of *NRGN* at 11q24.2. In functional magnetic resonance imaging (fMRI) studies, individuals carrying the risk variant showed significantly decreased activation in the hippocampus during contextual learning^18^, and widespread cortical thinning and thalamic shape abnormalities^19^. Moreover, a recent interactome analysis of another schizophrenia risk gene, *ZNF804A*, identified *NRGN* as one of its significant targets^20^.

Using the mouse model, our group recently reported that the translation of Ng is rapidly increased in response to neuronal activity, and this up-regulation is required for contextual memory formation^21^. Cognitive impairment is a core feature of the pathophysiology of schizophrenia, and reduced Ng immunoreactivity was observed in the prefrontal cortex regions of post-mortem brain tissues from schizophrenia patients^22^, suggesting that dysregulated Ng expression may contribute to the cognitive impairment in schizophrenia. Dysregulation of Ng expression has also been observed in Alzheimer’s disease^23, 24^, and the deletion of a chromosomal region containing the Ng gene causes a rare genetic disorder with symptoms of mental retardation, known as Jacobsen syndrome^25^. Ng levels are also dynamically regulated during development under different environmental and behavioral states^26-28^. Together, these evidences suggest that Ng is involved in cognitive impairment associated with neurodegenerative, neuropsychiatric and neurodevelopmental diseases.

Ng belongs to a family of small neuronal proteins, called calpacitins, which binds to a calcium-free form of calmodulin (CaM) via the IQ (isoleucine and glutamine-containing) domain. Among the calpacitins, Ng is uniquely expressed at high levels in the soma, dendrite and the postsynaptic compartment of the principal neurons in the cerebral cortex, hippocampus^26, 27^ and other brain regions important for experience-dependent plasticity including striatum and amygdala (Allen Brain Atlas)^29, 30^. CaM is released from Ng upon an increase in intracellular Ca^2+^ concentrations, and it is thought that Ng levels titrate the availability of CaM in the postsynaptic compartment of excitatory synpases in principal neurons^31, 32^.

CaM is a key signal transducer that detects the increase in cytosolic Ca^2+^ levels, which mediates Ca^2+^/CaM-dependent signaling events. The relative activation of Ca^2+^/CaM-dependent protein kinase II (CaMKII) and protein phosphatase 2B (PP2B, also known as calcineurin) at the postsynaptic compartment is considered to determine the direction of long-term potentiation (LTP), the cellular basis of learning and memory (Supplementary Fig. 1a)^33-41^. This view is in accordance with the push-pull mechanism that has been proposed for controlling the directionality and efficacy of synaptic plasticity^42-44^. Although both CaMKII and PP2B enzymes are activated by Ca^2+^/CaM complex, PP2B is preferentially activated when the amount of CaM is limited due to its much higher binding affinity for the Ca^2+^/CaM complex. In contrast, CaMKII is much more abundant than PP2B in the postsynaptic compartment, and thus CaMKII activity becomes dominant when the amount of Ca^2+^/CaM complex is sufficient^45-47^. Therefore, the expression of synaptic plasticity is highly sensitive to a change in CaM availability, and it was hypothesized that Ng controls synaptic plasticity by regulating CaM availability and Ca^2+^/CaM dynamics at the central excitatory synapses (Fig. 1a)^48, 49^. Consistent with the hypothesis, cortical neurons lacking Ng exhibit altered Ca^2+^ dynamics^50, 51^. However, previous studies with independently generated Ng knockout mice showed the contradicting effects in hippocampal LTP^50, 52, 53^, and results in young organotypic hippocampal slice cultures showed that elevated Ng levels occlude LTP under a standard pairing induction protocol, but facilitate LTP under a weaker pairing protocol^54, 55^. These data suggest that depending on the age, potentially genetic background and neural activity, Ng regulates synaptic plasticity differently. It is likely that a signaling network, rather than a single node in a particular pathway accounts for the mechanisms of LTP expression^56-62^, Moreover, recent genetic studies have revealed strong associations of genes involved in synaptic plasticity with neuropsychiatric and neurodevelopmental disorders, suggesting an intricate molecular interplay tuning synaptic plasticity critical for learning and memory^6, 63, 64^. It is therefore important to control the manipulation at defined developmental stage to be able to examine the scope and impact of Ng- dependent regulation on Ca^2+^/CaM-mediated signaling cascades and downstream targets and gain a comprehensive view of the molecular processes involved in synaptic plasticity.

**Fig. 1.**
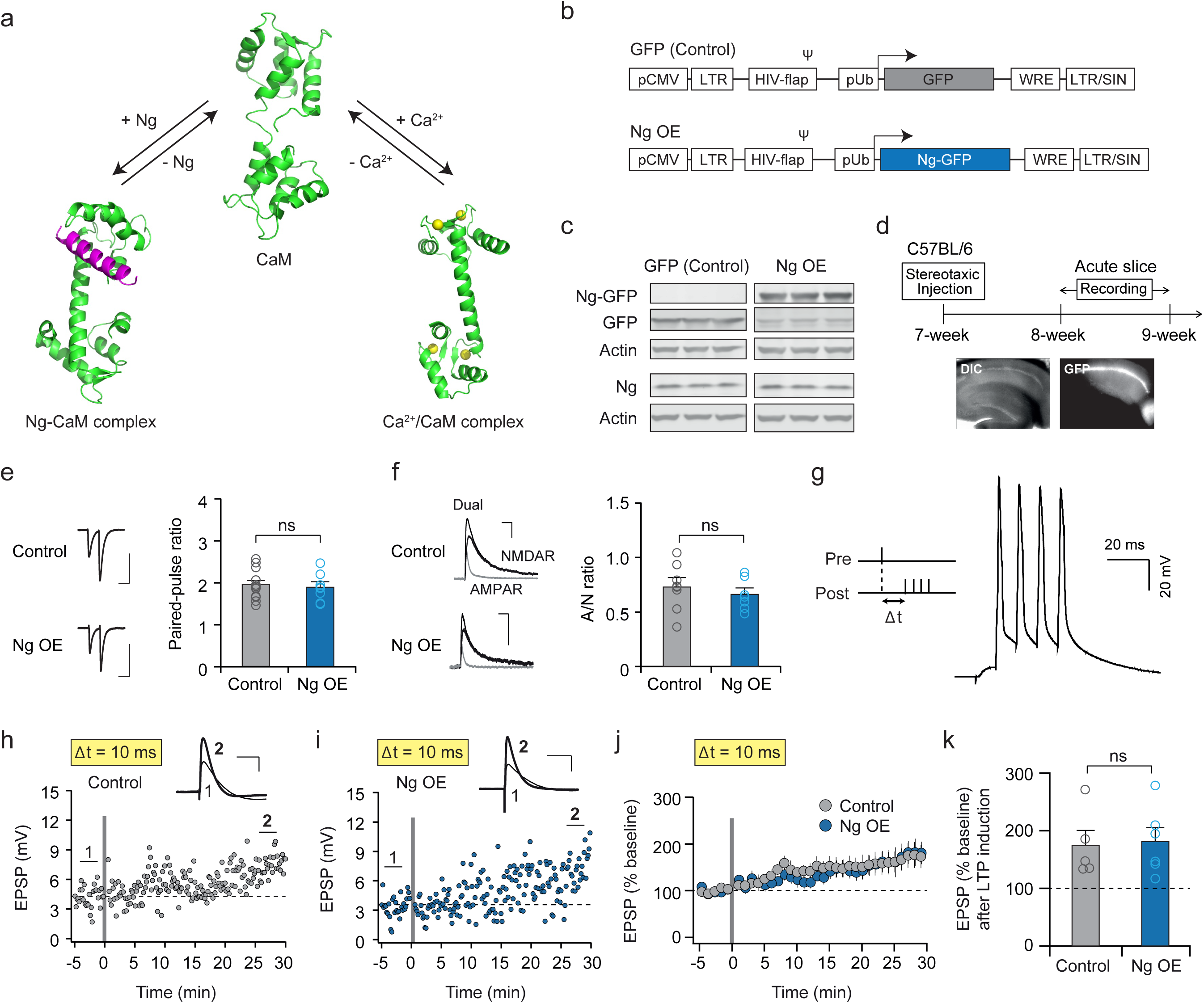
Ng overexpression does not affect basal synaptic transmission and pairing-induced LTP with 10-ms interval. (**a**) Ng binds to CaM and regulates the amount of CaM available for Ca^2+^ binding. Purple: Ng (only the CaM-binding IQ motif of Ng is depicted), Green: CaM, Yellow spheres: Ca^2+^ ions (PDB: CaM, 1CFD; Ca^2+^/CaM complex, 3CLN; Ng-CaM complex, 4E50). (**b**) Diagram of a lentivirus vector for Ng overexpression (Ng OE). LTR, long terminal repeat; Ψ, packing signal; Flap, flap element from HIV-1; pH1, H1 promoter; pUb, ubiquitin promoter; WRE, woodchuck hepatitis virus posttranscriptional regulatory element. (**c**) Immunoblot of cortical neuron culture infected with the Ng OE or GFP only (control) lentivirus shows effective overexpression of exogenous Ng-GFP fusion protein or GFP, respectively. (**d**) Experimental timeline for whole-cell patch clamp recordings is shown in the top panel. Bottom panel: The DIC and epifluorescence images show robust and exclusive expression of a lentiviral construct in the hippocampal CA1 region. (**e**) Comparison of paired-pulse ratio at 50-ms interval recorded from control and Ng OE neurons. Upper panel: average traces from control and Ng OE cells (scale bars, 100 pA, 50 ms). Bottom panel: collective data of paired-pulse ratio in control (n=12, 1.94 ± 0.10) and Ng OE (n=7, 1.89 ± 0.13) cells. The paired-pulse ratio values from individual cells are shown as small open circles. The average values are shown as filled circles with SEM (n.s.; not significant, t-test). (**f**) Comparison of NMDAR-EPSC to AMPAR-EPSC ratio in control and Ng OE neurons. Evoked EPSCs were measured in CA1 neurons following Schaffer collateral stimulation. Left panel: superimposed representative EPSC traces of dual components (compound EPSC of AMPAR and NMDAR), NMDAR-EPSC and AMPAR-EPSC measured at +40 mV. AMPAR-EPSC was obtained by application of D-APV, and NMDAR-EPSC was calculated by subtracting AMPAR- EPSC from dual components (scale bars, 50 pA, 50 ms). Right panel: collective data of the ratio of peak AMPAR-EPSC to NMDAR-EPSC in control (n=7, 0.73 ± 0.08) and Ng OE (n=6, 0.66 ± 0.06) cells. AMPAR/NMDAR ratio values from individual cells are shown as small open circles. The average values are shown as filled circles with SEM (n.s.; not significant, t-test). (**g**) Left panel: spike-timing-dependent plasticity was induced by 100 pairings of presynaptic and postsynaptic stimulations at 5 Hz. Each pairing consisted of stimulation at Schaeffer collaterals followed by four action potentials given at 100 Hz at various positive time intervals. An example of current clamp recording from a CA1 neuron during the pairing is shown in the right panel. (**h, i**) Sample recordings of STDP at 10-ms pairing interval from an uninfected control cell and a cell infected with Ng OE. Downward arrows indicate the timing of STDP induction. Traces show averaged EPSPs indicated with 1 and **2** (scale bars, 2 mV, 50 ms). (**j**) Averaged summary graphs of STDP at 10-ms interval in uninfected control (n=5) and Ng OE (n=6) cells. Each circle represents mean ± SEM. (**k**) Collective data of STDP at 10-ms interval in control (n=5, 182.4 ± 19.5 %) and Ng OE (n=6, 190.6 ± 17.8%) cells. EPSP after LTP induction (% baseline) values from individual cells are shown as small filled or open circles. The average values are shown as large filled or open circles with SEM (n.s.; not significant, t-test).

In this study, we used lentiviral-mediated gene transfer to manipulate Ng levels in CA1 neurons in adult brains, and sought to determine the influence of altered Ng levels on synaptic plasticity in the hippocampus, and found that Ng bidirectionally regulates spike-timing-dependent LTP at Schaffer collateral (SC) to CA1 synapses. To unbiasedly identify the underlying molecular mechanism, we applied quantitative phosphoproteomic analysis^65-67^ (Supplementary Fig. 1b), to reveal the molecular networks underlying the bidirectional control of LTP. That reduced Ng levels caused a significant shift in the phosphoproteome of postsynaptic density proteins, highlighting autism- and schizophrenia-associated gene targets. With further functional validation, our results suggest that altered Ng expression observed in neurodegenerative and neuropsychiatric diseases negatively affects the phosphorylation status of neuronal proteins including the NMDAR subunit Grin2A, by tuning synaptic PP2B activity, hence the induction and expression of synaptic plasticity, as an underlying cellular pathomechanism for the cognitive deficits in these diseases.

## Results

### Ng overexpression facilitates spike-timing-dependent LTP via direct interaction with CaM

To elucidate the role of Ng in synaptic plasticity, we used lentivirus-mediated manipulation of Ng levels in mouse hippocampal CA1 neurons, which allows post-developmental, postsynaptic neuron-specific manipulation and prevents potential complications resulting from developmental compensation. For Ng overexpression (Ng OE), a lentiviral construct to overexpress Ng was created using an ubiquitin promoter to drive the expression of Ng fused to GFP (Ng OE, Fig. 1b), and the robust expression of Ng-GFP fusion protein was confirmed by western blot (Fig. 1c). To manipulate the expression levels of Ng in CA1 neurons *in vivo*, we injected a concentrated lentivirus into the hippocampal CA1 region of 7-week-old C57BL/6 male mice by stereotaxic surgery. Acute hippocampal slices were prepared 5-9 days after the injection for electrophysiology recordings, and the lentiviral infection in the CA1 area was confirmed by GFP fluorescence (Fig. 1d). To examine the effect of Ng OE on the basal synaptic transmission, paired-pulse ratio (PPR) and AMPAR/NMDAR excitatory postsynaptic currents ratio (A/N ratio) at SC-CA1 synapses were recorded from uninfected cells (control) and infected cells from the same animals. Both PPR and A/N ratios were not significantly different between uninfected and infected neurons (Fig. 1e, f), indicating that increased levels of Ng in CA1 neurons do not alter presynaptic release probability and relative basal synaptic transmission at hippocampal SC-CA1 synapses under the basal condition.

The effect of elevated Ng levels on synaptic plasticity was examined using the spike-timing-dependent plasticity (STDP) protocol for LTP induction^68^. An individual pairing consisted of a presynaptic stimulation followed by a train of four action potentials repeated 100 times at 5 Hz (Fig. 1g). The relative timing between pre- and postsynaptic stimulations is a critical factor controlling plasticity for stimulated synapses^69, 70^. When the pairing was performed at a 10-ms interval, both uninfected and infected neurons expressed robust LTP with a similar degree (Fig. 1h-k), indicating that increased Ng levels exert no additional effects on the magnitude of LTP when the induction protocol with the 10-ms pairing interval is used, and endogenous Ng levels are sufficient to support the expression of STDP-LTP.

We then examined the effect of Ng OE on STDP-LTP under a weaker induction condition, and a prolonged pairing interval is known to drive less cooperated activity between presynaptic release and postsynaptic membrane depolarization. When the pairing interval was increased to 20 ms, the pairing protocol no longer induced LTP in control cells (Fig 2a, g, h), whereas neurons with Ng OE showed robust LTP expression (Fig. 2b, g, h). This result indicates that increased Ng levels allow the expression of STDP-LTP with the prolonged pairing interval, and Ng facilitates the induction of STDP-LTP by broadening the temporal association window.

**Fig. 2.**
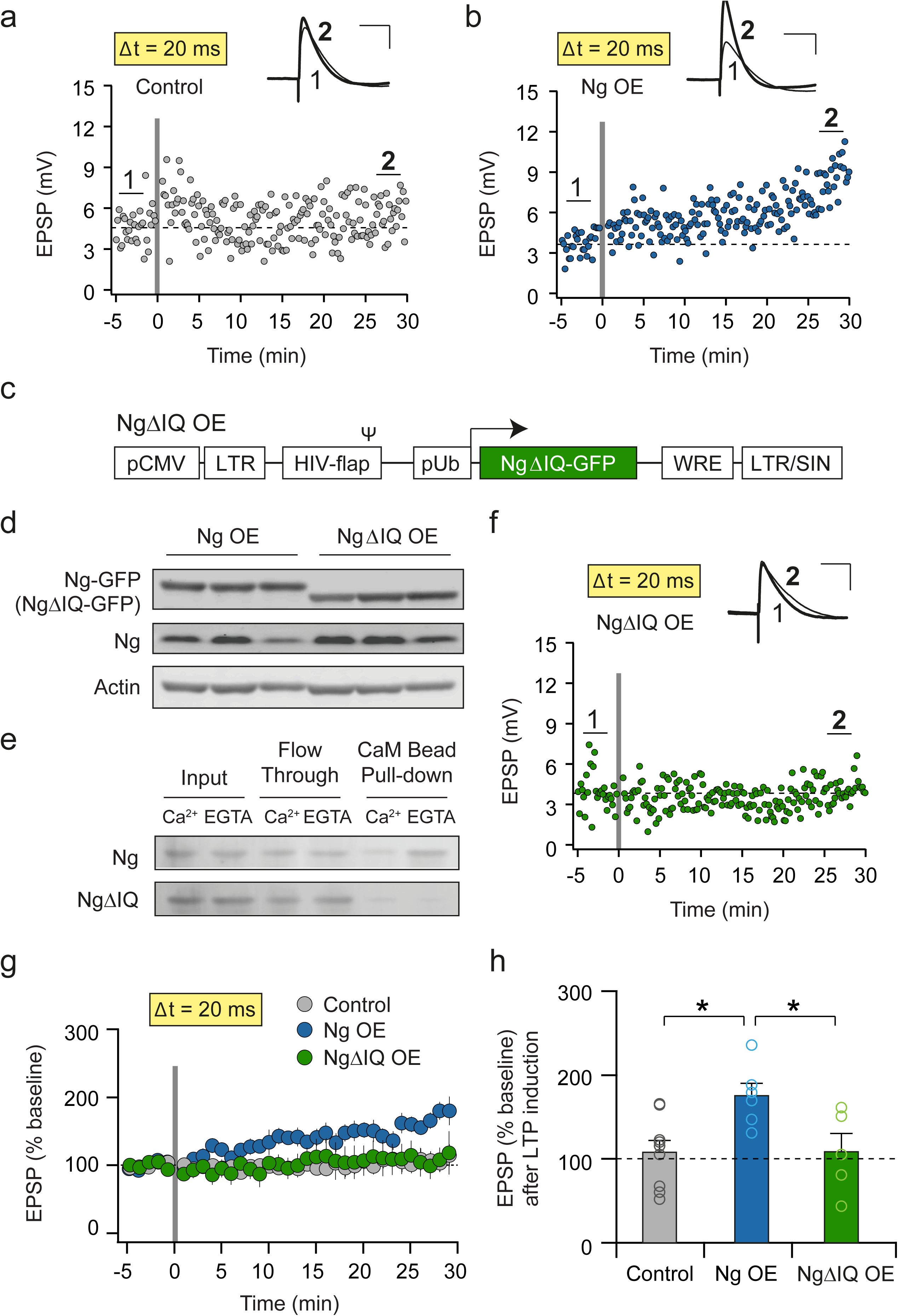
Ng overexpression facilitates the induction of LTP with 20-ms pairing interval. (**a, b**) Sample recordings of STDP at 20-ms pairing interval from an uninfected control cell and a cell infected with Ng OE. Downward arrows indicate the timing of STDP induction. Traces show averaged EPSPs indicated with 1 and **2** (scale bars, 2 mV, 50 ms). (**c**) Diagram of a lentivirus vector for a Ng mutant lacking the CaM-binding IQ motif (NgΔIQ). (**d**) Immunoblot of cortical neuron culture infected with the Ng OE or the Ng deletion mutant lentivirus shows effective overexpression of indicated constructs. The deletion mutant runs a bit smaller compared to the wildtype Ng, as expected. (**e**) Binding of Ng to CaM was examined by pull-down assay in the presence of 2 mM Ca^2+^ or 2 mM EGTA. Immunoblot of total cell lysate (input), flow through, and proteins bound to the CaM beads (CaM bead pull-down) samples probed with an antibody against Ng C-terminal. (**f**) A sample recording of STDP at 20-ms pairing interval from a cell infected with NgΔIQ. The downward arrow indicates the timing of STDP induction. Traces show averaged EPSPs indicated with 1 and **2** (scale bars, 2 mV, 50 ms). (**g**) Averaged summary graphs of STDP at 20-ms interval in uninfected control (n=9), Ng OE (n=6) and NgΔIQ (n=5) cells. Each circle represents mean ± SEM. (**h**) Collective data of STDP at 20-ms interval in control (n=9, 104.2 ± 10.4%), Ng OE (n=6, 189.8 ± 23.2%) and NgΔIQ (n=5, 106.5 ± 19.8%) cells. EPSP after LTP induction (% baseline) values from individual cells are shown as small filled or open circles. The average values are shown as large filled or open circles with SEM (\**p*<0.05, One-way ANOVA and Tukey’s multiple comparison test).

To test whether this facilitative effect is mediated by the direct interaction between Ng and CaM, a mutant form of Ng was created in which the CaM-binding IQ motif is deleted (Fig. 2c, d, NgΔIQ). HEK cells were transiently transfected with a construct expressing either wildtype Ng or NgΔIQ, and the cell lysates were incubated with beads coated with purified CaM to confirm the specific and Ca^2+^-dependent interaction between CaM and Ng. In agreement with previous studies, wildtype Ng was preferentially bound to CaM under the low Ca^2+^ condition (EGTA), but NgΔIQ did not interact with CaM regardless of a change in Ca^2+^ concentrations (Fig. 2e), confirming that Ng directly interacts with CaM through the IQ motif. As expected, when the NgΔIQ mutant was overexpressed in CA1 neurons, the induction protocol with a 20-ms interval was not able to trigger LTP (Fig. 2f-h), suggesting that the interaction with CaM is critical for the facilitative effect of Ng on the induction of LTP.

### Ng knockdown abolishes the induction of STDP-LTP at SC-CA1 synapses

We next asked whether the Ng-dependent regulation of STDP-LTP is bidirectional. For Ng knockdown (Ng KD), a lentiviral vector with dual promoters was constructed, in which the H1 promoter drives the expression of shRNA targeting endogenous Ng mRNAs and the ubiquitin promoter drives the simultaneous expression of GFP as an infection marker (Fig. 3a, Ng KD). The knockdown of Ng was highly effective 7-10 days after viral infection when tested by western blot in dissociated cortical neuron culture (Fig. 3b, control bands are the same as those in Fig. 1c, as GFP-infected cultures were used as controls for both conditions in one preparation). PPR and A/N ratio were recorded from both uninfected control cells and infected cells from the same animals, and Ng KD did not significantly affect PPR and A/N ratio at SC-CA1 synapses (Fig. 3c, d), indicating that presynaptic release probability and basal synaptic transmission remain intact at SC-CA1 synapses.

**Fig. 3.**
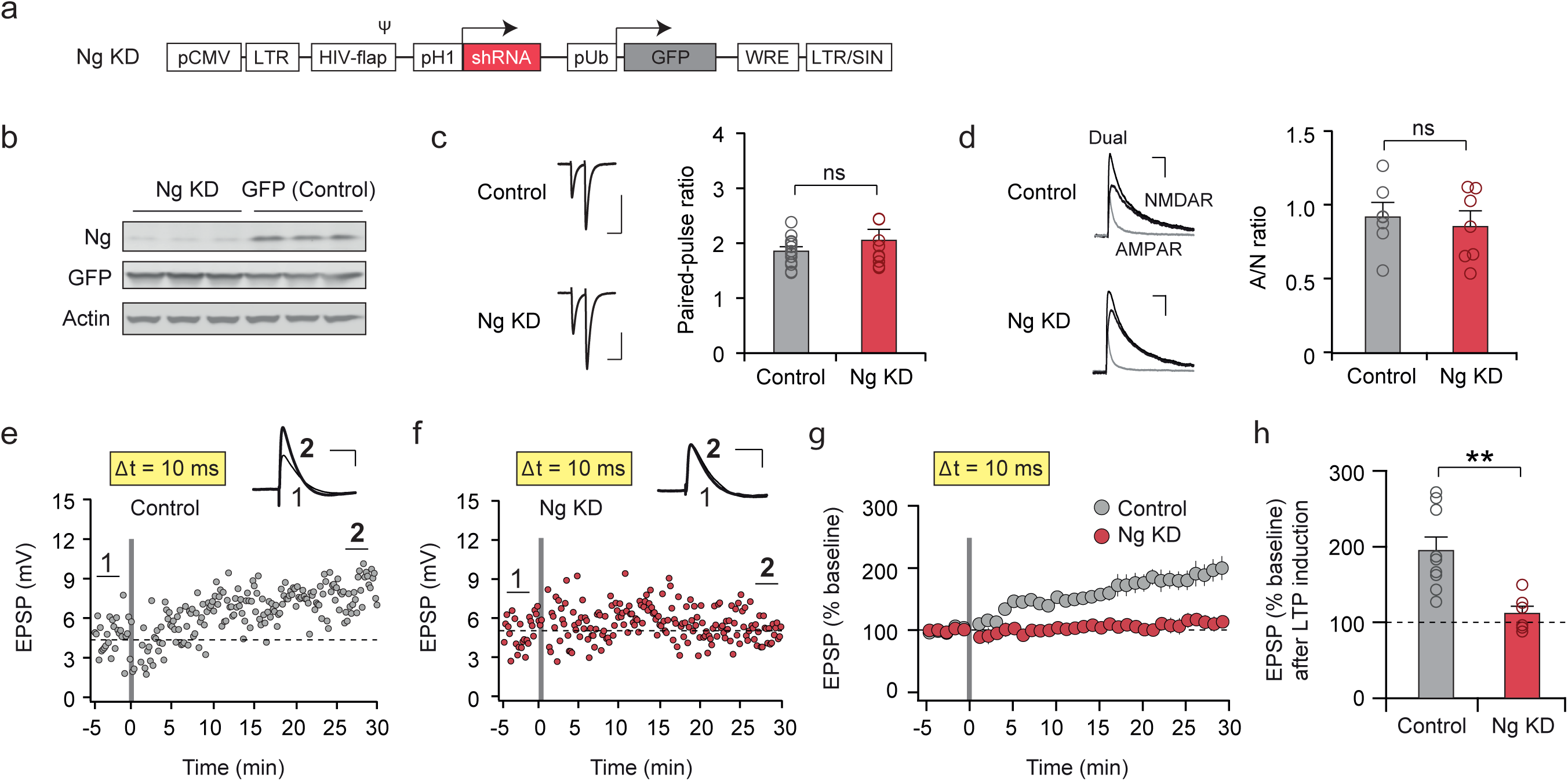
Ng knockdown abolishes the induction of LTP at SC-CA1 synapses. (**a**) Diagram of a lentivirus vector for Ng knockdown (Ng KD). LTR, long terminal repeat; Ψ, packing signal; Flap, flap element from HIV-1; pH1, H1 promoter; pUb, ubiquitin promoter; WRE, woodchuck hepatitis virus posttranscriptional regulatory element. (**b**) Immunoblot of cortical neuron culture infected with the Ng KD or GFP only (control) lentivirus shows effective knockdown of endogenous Ng. (**c**) Comparison of paired-pulse ratio at 50-ms interval recorded from control and Ng KD neurons. Upper panel: average traces from control and Ng KD cells (scale bars, 100 pA, 50 ms). Bottom panel: collective data of paired-pulse ratio in control (n=12, 1.85 ± 0.08) and Ng KD (n=9, 2.05 ± 0.20) cells. The paired-pulse ratio values from individual cells are shown as small open circles. The average values are shown as filled circles with SEM (n.s.; not significant, t-test). (**d**) Comparison of NMDAR-EPSC to AMPAR-EPSC ratio in control and Ng KD neurons. Evoked EPSCs were measured in CA1 neurons following Schaffer collateral stimulation. Left panel: superimposed representative EPSC traces of dual components (compound EPSC of AMPAR and NMDAR), NMDAR-EPSC and AMPAR-EPSC measured at +40 mV. AMPAR- EPSC was obtained by application of D-APV, and NMDAR-EPSC was calculated by subtracting AMPAR-EPSC from dual components (scale bars, 50 pA, 50 ms). Right panel: collective data of the ratio of peak AMPAR-EPSC to NMDAR-EPSC in control (n=6, 0.91 ± 0.10) and Ng KD (n=6, 0.85 ± 0.11) cells. AMPAR/NMDAR ratio values from individual cells are shown as small open circles. The average values are shown as filled circles with SEM (n.s.; not significant, t-test). (**e, f**) Sample recordings of STDP at 10-ms pairing interval from an uninfected control cell and a cell infected with Ng KD. Downward arrows indicate the timing of STDP induction. Traces show averaged EPSPs indicated with 1 and **2** (scale bars, 2 mV, 50 ms). (**g**) Averaged summary graphs of STDP at 10-ms interval in uninfected control (n=9) and Ng KD (n=6) cells. Each circle represents mean ± SEM. (**h**) Collective data of STDP at 10-ms interval in control (n=9, 196.3 ± 15.2 %) and Ng KD (n=6, 105.1 ± 10.8 %) cells. EPSP after LTP induction (% baseline) values from individual cells are shown as small filled or open circles. The average values are shown as large filled or open circles with SEM (\*\**p*<0.01, t-test).

The effect of decreased Ng levels on STDP-LTP was examined using the pairing protocol with 10-ms pairing interval, and a robust LTP was expressed in non-infected control neurons (Fig. 3e, g, h). Conversely, the induction of LTP was completely abolished in neurons infected with Ng KD (Fig. 3f-h), indicating the essential role of Ng for STDP-LTP expression.

### Decreased Ng levels cause a significant shift in the postsynaptic phosphoproteome, including hypo-phosphorylation of NMDAR subunit Grin2A

Given the role of Ng in regulating CaM availability as well as Ca^2+^/CaM dynamics, we hypothesized that Ng KD influences the activation of Ca^2+^/CaM-dependent kinases and phosphatases, thereby leading to a global change in protein phosphorylation as well as altered synaptic plasticity. To examine the overall changes in the phosphoproteome, we enriched phosphopeptides using immobilized metal affinity chromatography (IMAC) from total cell lysates prepared from dissociated neuronal cultures infected with either GFP or Ng KD lentiviruses at DIV 7 and collected at DIV 17. Both total proteome and phosphoproteome were analyzed using quantitative mass spectrometry with isobaric labeling of peptides as previously described (Fig. 4a)^71, 72^. The phosphoproteome data were normalized to the total proteome when the proteins were confidently identified in the total proteome dataset (Supplementary Table 1). Nearly 30,000 phosphorylation sites (p-sites) comprising of 5,485 proteins identified in the proteome were analyzed. 4,744 (∼16%) of these p-sites derived from 2,413 proteins exhibited a significant change in their phosphorylation status compared to the control (Fig. 4b, FDR ≤ 0.05). These data show that decreasing Ng levels in neurons induced a significant shift in the phosphoproteome landscape.

**Fig. 4.**
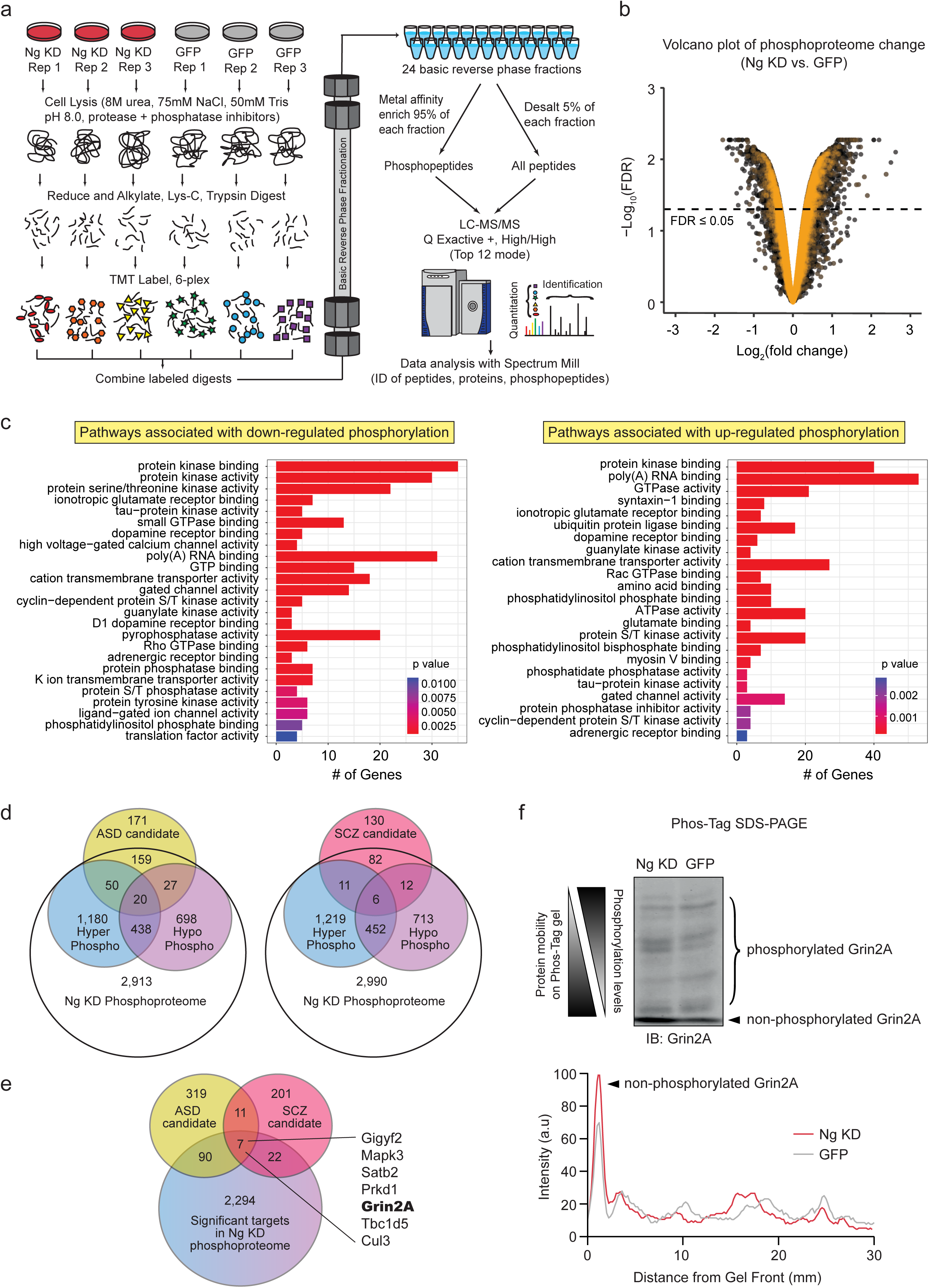
Knockdown of Ng causes significant shifts in neuronal phosphoproteome. (**a**) Proteomic and phosphoproteomic workflow for the Ng KD experiment. Ng KD and the respective GFP controls were grown in triplicate (n=6), collected, and lysed in 8M urea with protease and phosphatase inhibitors. The denatured protein was reduced, and alkylated, and double digested with both Lys-C and Trypsin overnight. The tryptic peptides were labeled with TMT- 6 reagent and the individual label incorporation was checked via LC-MS/MS. The labeled digests were combined and basic reverse phase fractionated into 24 fractions. From each fraction, 5% of the total volume was remove for proteomic analysis while the remaining 95% was used for phosphopeptide enrichment. The proteome and the phosphoproteome data was acquired on a Q-Exactive + mass spectrometer. Peptide spectrum matching and protein identification was performed using Spectrum Mill. (**b**) Volcano plots comparing the individual phosphoproteome phosphorylation sites of the Ng KD experiment. The −log_10_ of the adjusted *p* value is plotted against the average log_2_ fold change for the phosphoproteome. The dotted line represents an adjusted *p* value of 0.05. Orange points represent identified post-synaptic density components as described by ^73^. (**c**) Left: The pathways highlighted with down-regulated phosphorylation of PSD targets in Ng KD. Right: The pathways highlighted with up-regulated phosphorylation of PSD targets in Ng KD. (**d**) Left: The overlap of significantly affected phosphorylated targets by Ng KD with the ASD- associated gene set. Right: The overlap of significantly affected phosphorylated targets by Ng KD with the schizophrenia-associated gene set. The upregulated phosphoproteome with Ng KD shown in blue; the downregulated phosphoproteome with Ng KD shown in purple; the ASD gene set shown in yellow; and the schizophrenia gene set shown pink. (**e**) The overlap of significantly affected phosphorylated targets by Ng KD with both the ASD- and schizophrenia-associated gene sets. (**f**) A differential phosphorylation of NMDAR subunit Grin2A was examined using a Phos-Tag SDS-PAGE, and Ng KD led to an increase in the fraction of non-phosphorylated Grin2A subunit.

Using the hypergeometric test, we found that differentially regulated p-sites in the phosphoproteome dataset were significantly over-represented in the set of known postsynaptic density (PSD) proteins (p<2×10^−11^, Supplementary Table 1)^73^. Specifically, 26% of the proteins with significantly down-regulated p-sites overlapped with the PSD proteomic dataset (∼29% of all significantly down-regulated p-sites), and 27% of the proteins with significantly up-regulated p-sites overlapped with the PSD proteomic dataset (∼33% of all significantly up-regulated p-sites) (Supplementary Table 1), indicating that decreasing Ng levels significantly shifted the phosphorylation state of postsynaptic components. To further determine which cellular functions are most directionally affected under Ng KD, we performed GO enrichment analysis^74, 75^, separately on the sets of up- and down-phosphorylated PSD proteins using the clusterProfiler R package^76^. Notably, pathways related with protein kinase binding were highlighted in both of the up- and down-regulated clusters. Also, ion transport and ion channel binding clusters were highlighted in the set with down-regulated phosphorylation (Fig. 4c).

Given the association of Ng with mental retardation and schizophrenia, and the convergence of the glutamatergic synaptic components in schizophrenia and autism spectrum disorders (ASDs), we questioned whether the changes in Ng levels influence the phosphorylation states of ASD- and schizophrenia-associated gene targets. In order to answer this question, we took the human ASD gene list from SFARI (https://gene.sfari.org/autdb/HG_Home.do) with category scores ≤ 4. Among the 460 genes included in the list, 427 were converted to mouse genes and compared with the list from our proteomics analysis (Fig. 4d, left). 256 (60%) of 427 were identified in the phosphoproteome data, and 37.9% of the gene targets were identified (97 out of 256) with significant changes in phosphorylation states with Ng KD. The list of genes with phosphorylation sites are shown in Supplementary Table 2, clustered using DAVID functional annotation^77, 78^. Notably, synaptic components and ion channels were highlighted, indicating the overlap of ASD targets and phosphoproteome changes induced by Ng KD.

In addition, we also took the candidate genes from the 108 Loci associated with schizophrenia^6^. Among 333 extracted human genes, 241 of them were converted into mouse genes. 111 (46%) of these were identified in the phosphoproteome, in which 29 (26%) were identified with significant changes in phosphorylation states with Ng KD (Fig. 4d, right; Supplementary Table 3). Importantly, among the proteins whose phosphorylation states were significantly altered by Ng KD, seven targets were associated with both ASD and schizophrenia (Fig. 4e), highlighting a potential convergence of the pathomechanisms of ASD and schizophrenia. Out of these identified targets, the NMDAR subunit Grin2A (also known as NR2A, GluN2A) was of particular interest, given its important role in conducting Ca^2+^ ions critical for NMDAR-dependent plasticity^79-81^. We identified several phosphorylation sites in the C-terminal region of Grin2A that were significantly affected by Ng KD. In particular, Grin2A S1384, a previously uncharacterized phosphorylation site, was hypo-phosphorylated by Ng KD. In addition, the phosphorylation site Grin2A S882/S890 was significantly hyper-phosphorylated. To further examine the overall change in phosphorylation status of Grin2A in the Ng KD condition, we used a Phos-Tag SDS-PAGE gel system, which separates a protein based on the degree of phosphorylation levels^82, 83^, and revealed that Ng KD shifted the phosphorylation pattern of Grin2A toward the hypo-phosphorylated state (migrating to the forefront of the Phos-Tag gel; Fig. 4f).

### C-terminal phosphorylation of Grin2A modulates NMDAR-mediated current kinetics

To investigate the functional significance of hypo-phosphorylation of Grin2A on NMDAR-mediated currents, we focused on the down-regulated phosphorylation sites of Grin2A identified from the phosphoproteome data. Specifically, the phosphorylation of Grin2A S1384 (Uniprot ID: P35436) was significantly decreased with Ng KD. In a separate sample set, the hypo-phosphorylation of Grin2A at S1384 was independently validated, and three additional sites S1198, S1201, S1204 also showed significant hypo-phosphorylation, implying the tolerance at certain phosphorylation sites. Given the profound hypo-phosphorylation pattern of Grin2A observed with a Phos-Tag gel, we generated Grin2A mutants in which the four phosphorylation sites were mutated to alanine (SA; phospho-deficient), or aspartic acid (SD; phospho-mimetic) residues to determine the role of the four serine residues in regulating NMDAR functions (Fig. 5a). Given that all four phosphorylation sites are positioned in the C-terminus, a C-terminal truncated mutant (-Ct) was also created as an additional control for C-terminus-mediated functions (Fig. 5a). The Grin2A mutants were co-expressed with GFP-fused Grin1 subunit separated by a self-cleaving P2A peptide in single-copy, isogenic, inducible HEK 293 cells (Fig. 5a, b). Stable cell lines were induced with doxycycline (Dox), and the expression of constructs was validated by qPCR, western blot, and live cell confocal microscopy (Fig. 5c; see also Supplementary Fig. 2a, b). Their responses to glutamate pulse were evaluated using high-throughput single-cell planar patch clamp with the SyncroPatch 384PE (Fig. 5d, left)^84^. Robust inward currents were elicited upon application of 10 μM glutamate in the presence of 30 μM glycine (Fig. 5d, right).

**Fig. 5.**
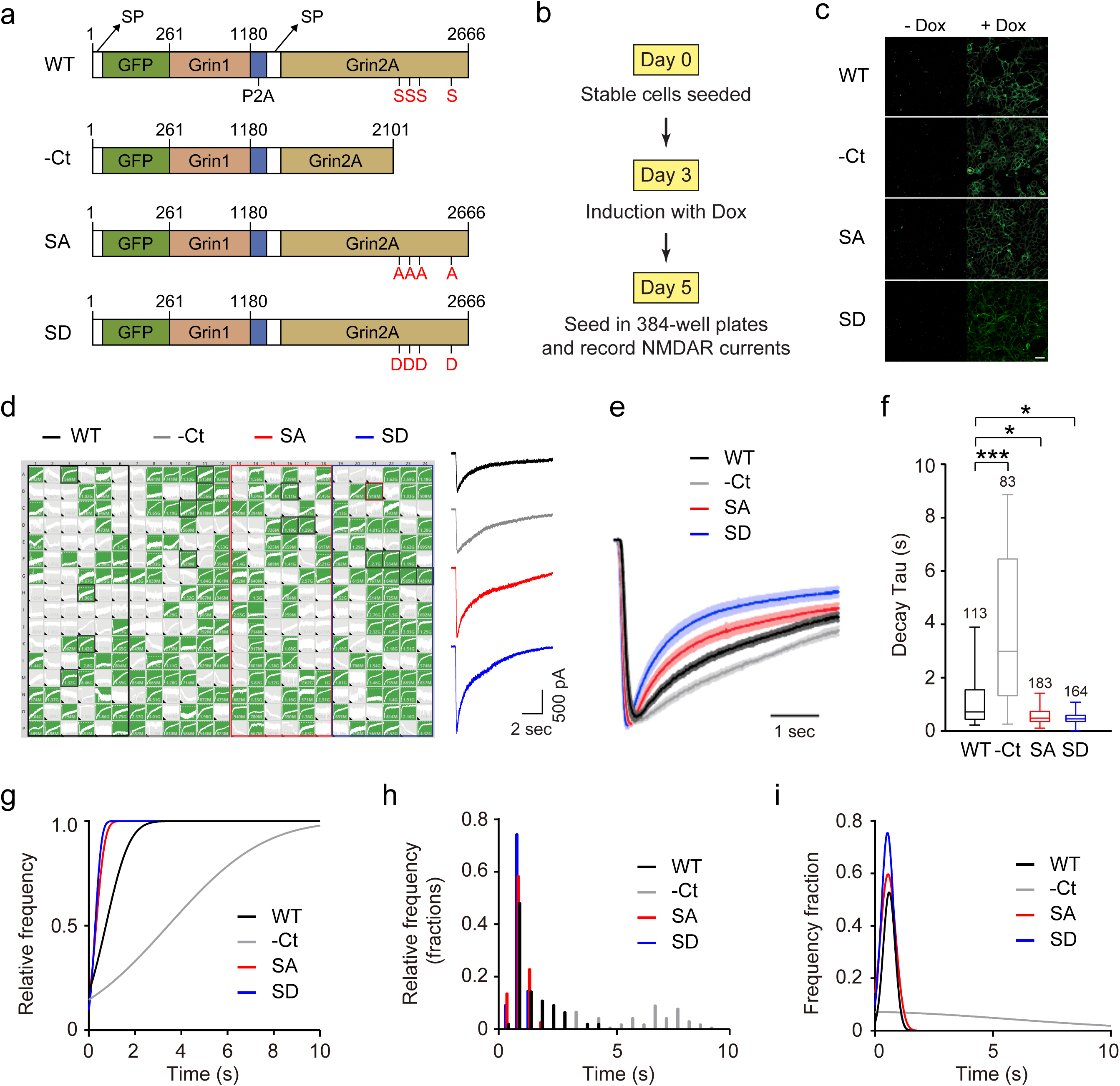
C-terminal phosphorylation of Grin2A modulates NMDAR-mediated current kinetics. (**a**) Design of wild-type (WT), C-terminus deletion (-Ct), serine to alanine (SA), and serine to glutamate (SD) isogenic, single-copy, doxycycline-inducible NMDAR expression constructs. The four phosphorylation sites selected for mutation are S1198, S1201, S1204, and S1384 in rGrin2A (Uniprot ID: P35436). (**b**) Experimental design for throughput analysis of NMDAR-mediated currents using a 384-well planar patch clamp electrophysiology system. (**c**) Live cell confocal images demonstrating the surface expression of NMDAR WT and mutants 48 hours post-induction with doxycycline, scale bar, 25 μm. (**d**) Left: Example of a 384-well (16 by 24) planar patch clamp recording. Right: Representative recordings of NMDAR-mediated currents using planar patch clamp. (**e**) Average traces of NMDAR currents with Grin2A WT and mutant normalized to peak current highlight differences in decay kinetics in –Ct, SA and SD mutants. Shaded bands represent SEM. (**f**) Box plots of decay Tau values of NMDAR currents recorded from the cell lines with Grin2A WT, and –Ct, SA and SD mutants. Data were compared via one-way ANOVA and significance was calculated with the Holmes-Sidak multi-comparisons test. \**p*<0.05, \*\*\**p*<0.001. (**g-i**) Gaussian fits of the cumulative distribution of decay kinetics (g), probability density histograms of decay kinetics (h) and its Gaussian fits (i) demonstrate Gaussian distributions for all experimental conditions except for -Ct.

Given that the C-terminal region of the Grin subunits regulates the NMDAR channel kinetics^85-87^ and a change in the phosphorylation state of Grin2A subunits influences the kinetics of synaptic NMDAR currents^88-91^, we hypothesized that the four serine sites in the C-terminus in Grin2A regulates NMDAR current kinetics, and compared the decay of NMDAR current from WT and mutants expressed in HEK 293 cells. SA and SD mutants exhibited significantly faster decay of NMDAR-mediated currents compared to WT and the –Ct mutant (Fig. 5e-i). The decay kinetics was not correlated with the amplitude of peak currents in all four mutants (Supplementary Fig. 2c, d), indicating that the difference among the mutants is not due to the amount of Ca^2+^ influx. This is further supported by the recording in the Ba^2+^ containing solution, in which the SA and SD mutants also exhibited significantly faster decay compared to WT and the –Ct mutant, regardless of the size of the peak currents (Supplementary Fig. 2e-j). Given that a significant amount of Grin2A subunit existed in the hypo-phosphorylated state in the Ng KD condition, it is likely that both SA and the SD mutants mimic the dephosphorylated state of Grin2A at these phosphorylation sites, which has been shown with other proteins^92-95^.

We also found that the rise kinetics of NMDAR-mediated currents was accelerated in – Ct, SA, and SD mutants compared to WT when recorded in the presence of Ca^2+^, and the rise kinetics was not correlated with the size of the peak current (Supplementary Fig. 3a-f). However, the difference in rise kinetics was absent in the Ba^2+^ recording condition (Supplementary Fig. 3g-k), suggesting that a Ca^2+^-dependent process also regulates channel activation via the phosphorylation of the Grin2A C-terminus.

Together, these data suggest that the phosphorylation state of Grin2A C-terminus regulates the kinetics of NMDAR currents, and the dephosphorylation of the four serine sites (S1198, 1201, 1204 and 1384) accelerates the current decay.

### Decreased Ng levels accelerates the decay of synaptic NMDAR currents by elevating PP2B activity

Given that the induction of STDP-LTP requires Ca^2+^ influx through NMDARs^96^, that phosphorylation of the Grin2A subunit was significantly decreased with Ng KD (Fig. 4), and that the Grin2A SA and SD mutants exhibited accelerated decay of NMDAR current (Fig. 5), we examined whether the kinetics of synaptically evoked NMDAR current is altered by reduced Ng levels in the hippocampus as a potential cause for the STDP-LTP deficit seen in the Ng KD condition (Fig. 3). The NMDAR-mediated currents recorded at SC-CA1 synapses were best fitted with a two-component exponential function^97^. While the slow component was not significantly different between the non-infected neurons and the neurons infected with Ng KD (data not shown), the fast component had a smaller decay time constant in Ng KD (Fig. 6a). The accelerated decay of synaptically evoked NMDAR-mediated currents observed in Ng KD is consistent with the case of SA and SD mutants in HEK 293 cells, suggesting that a decrease in Ng expression makes Ca^2+^ influx through NMDARs at synapses more transient by the de-phosphorylation of Grin2A.

**Fig. 6.**
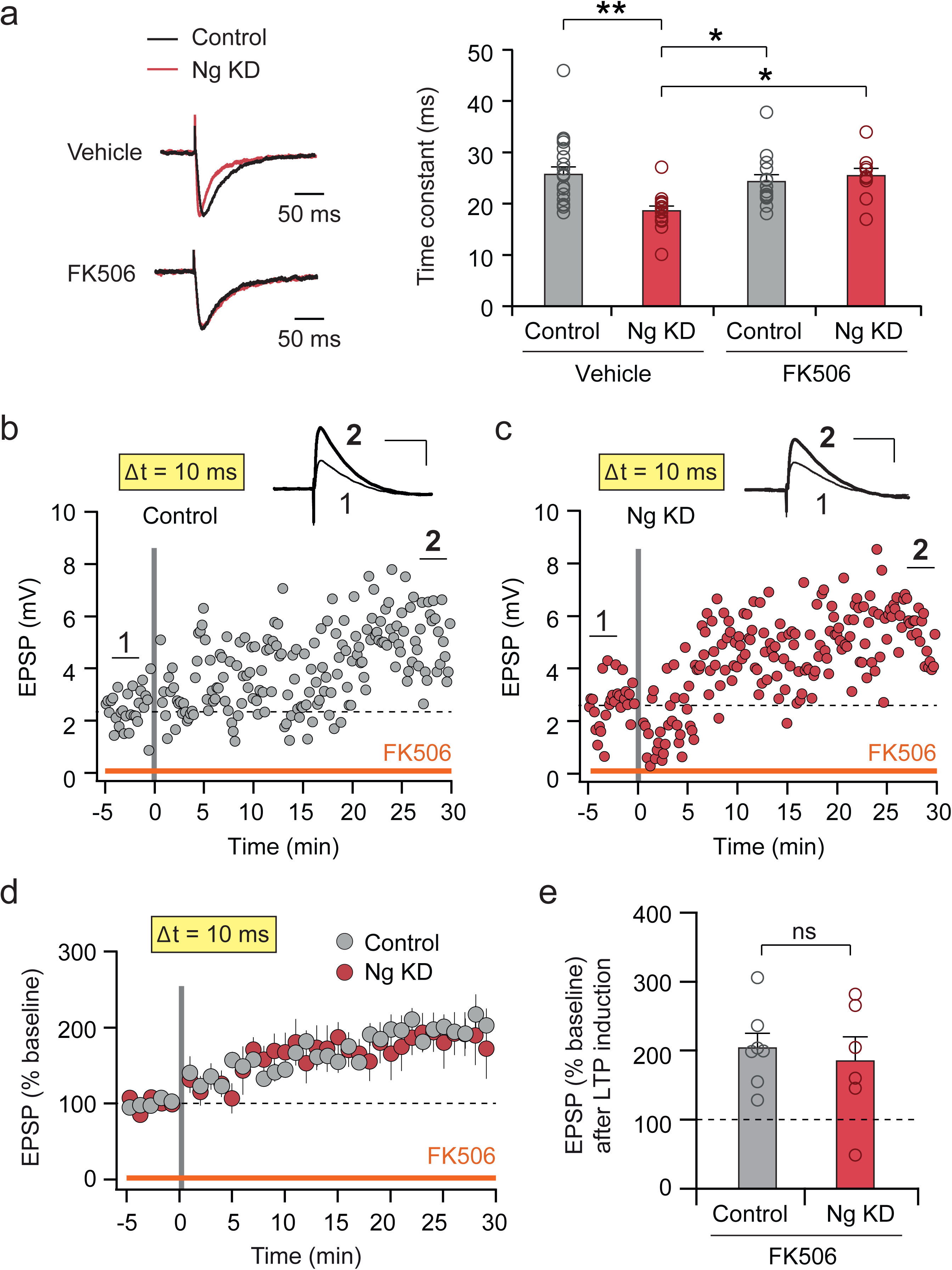
Ng knockdown accelerates the decay of NMDAR-mediated synaptic currents in SC-CA1 synapses by increasing PP2B activity. (**a**) Comparison of NMDAR-mediated calcium currents recorded from control and Ng KD neurons. Left panel: average traces of NMDAR currents from control (thin line) and Ng KD (thick line) cells recorded in vehicle only (upper) or in the presence of 1 μM FK506 (bottom). NMDAR currents were fitted with a two-exponential decay function, and the slow component were not significantly different across the four conditions. Right panel: collective data of the fast component of NMDAR currents measured from control (n = 21, 25.70 ± 1.44 ms) and Ng KD (n = 15, 18.63 ± 0.92 ms) cells in vehicle and from control (n = 14, 24.31 ± 1.34 ms) and Ng KD (n = 10, 25.48 ± 1.41 ms) cells in FK506. The fast component of exponential decay values from individual cells are shown as small open circles. The average values are shown as filled circles with SEM (\*\**p*<0.01, \**p*<0.05, Two-way ANOVA and Tukey’s multiple comparison test). (**b, c**) Sample recordings of STDP at 10-ms pairing interval from an uninfected control cell and a cell infected with Ng KD in the presence of FK506. Downward arrows indicate the timing of STDP induction. Traces show averaged EPSPs indicated with 1 and **2** (scale bars, 2 mV, 50 ms). (**d**) Averaged summary graphs of STDP at 10-ms interval in uninfected control (n=7) and Ng KD (n=6) cells in the presence of FK506. Each circle represents mean ± SEM. (**e**) Collective data of STDP at 10-ms interval in control (n=7, 203.6 ± 17.8 %) and Ng KD (n=6, 165.1 ± 35.8%) cells in the presence of FK506. EPSP after LTP induction (% baseline) values from individual cells are shown as small filled or open circles. The average values are shown as large filled or open circles with SEM (n.s.; not significant, t-test).

With decreased Ng levels, more CaM (and thus more Ca^2+^/CaM complex) become available for Ca^2+^ binding under a resting condition, which may lead to elevated activation of Ca^2+^/CaM-dependent phosphatase PP2B due to its high affinity toward the Ca^2+^/CaM complex^98^. Therefore, we tested whether suppressing PP2B activity could rescue the accelerated decay of synaptic NMDAR-mediated currents using FK506, a PP2B antagonist. The application of FK506 had no effect on the decay kinetics of NMDAR currents in the control neurons, but it rescued the accelerated decay in Ng KD to the control level (Fig. 6a). These results suggest that the basal synaptic PP2B activity in control neurons, if any, does not affect the kinetics of NMDAR currents. In Ng KD, however, decreased Ng levels increase synaptic PP2B activity, which in turn dephosphorylates the Grin2A subunit at synapses and accelerates the decay of NMDAR-mediated currents^88, 89^.

PP2B plays an important role in regulating synaptic plasticity and memory formation^42, 99, 100^, and the elevated PP2B activity could be responsible for the impaired LTP in Ng KD. To test this idea, the effect of FK506 treatment on the STDP-LTP was examined with a pairing protocol of a 10-ms interval. Notably, FK506 rescued LTP in Ng KD (Fig. 6c-e) without affecting the magnitude of LTP in the control cells (Fig. 6b, d, e). Taken together, these results suggest that a decrease in Ng expression elevates PP2B activity, which dephosphorylates synaptic Grin2A subunits and accelerates the decay of synaptic NMDAR currents, thereby leading to the impairment of LTP.

### Ng overexpression and decreased PP2B activity converge on the facilitation of STDP-LTP

Given the role of Ng in regulating the availability of CaM, and the effect of Ng KD on phosphoproteome and synaptic PP2B activity, we examined whether the facilitation of LTP by Ng OE also involves the modulation of PP2B activity. As opposed to the case of Ng KD in which more CaM become available to boost basal PP2B activity, Ng OE is expected to suppress the formation of Ca^2+^/CaM complexes by sequestering more CaM and thus inhibit basal PP2B activity. If decreased PP2B activity caused by Ng OE plays a crucial role in the facilitation of STDP-LTP, then blocking PP2B activity is likely to mimic the facilitative effect on STDP-LTP. Interestingly, bath application of FK506 enabled control cells to express robust STDP-LTP with the 20-ms pairing protocol which used to be a sub-threshold protocol in control cells (Fig. 7a, b, e-g), suggesting that basal PP2B activity inhibits control cells from expressing LTP under the 20-ms pairing protocol.

**Fig. 7.**
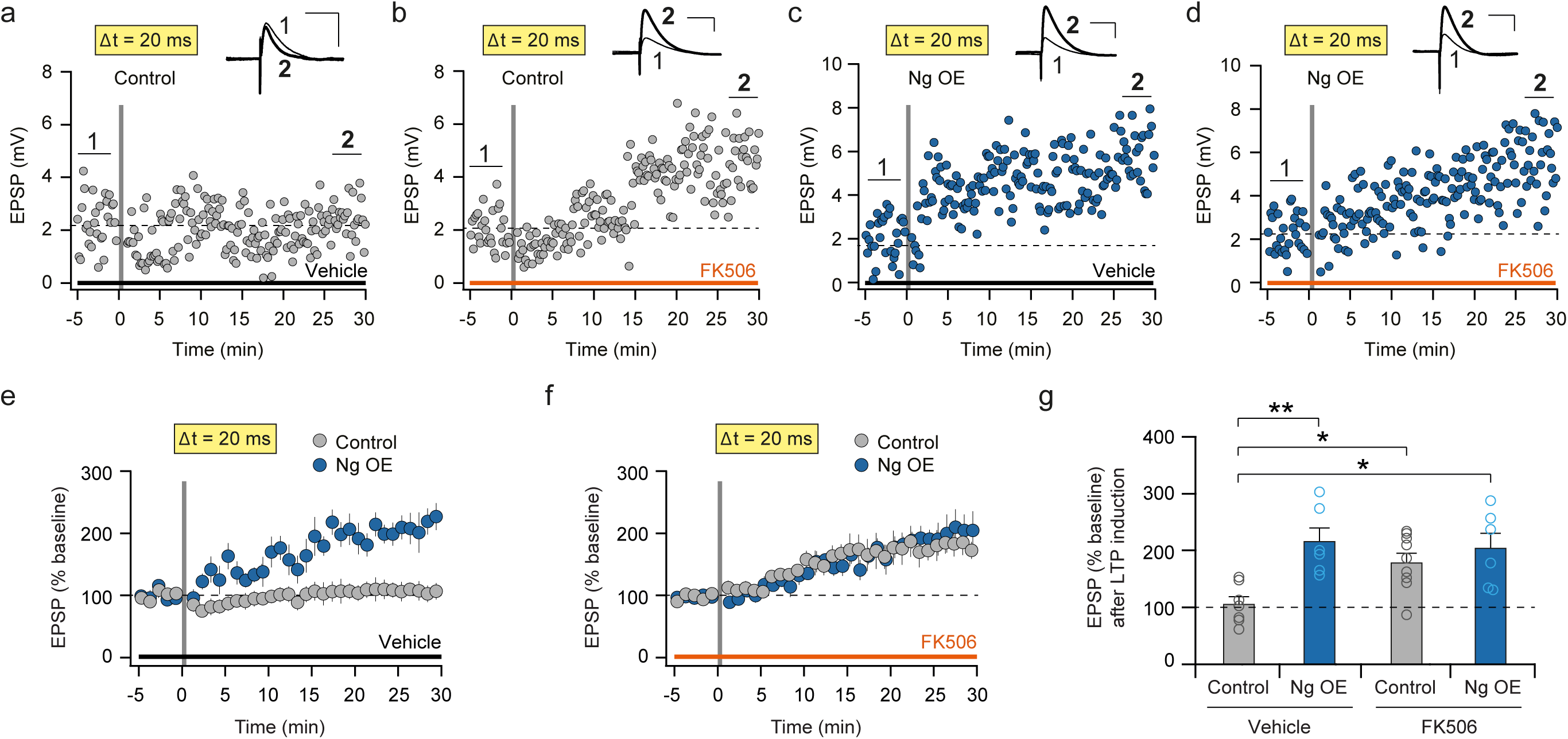
Ng overexpression facilitates LTP by suppressing PP2B activity. (**a-d**) Sample recordings of STDP at 20-ms pairing interval from an uninfected control cell in vehicle only, a control cell in the presence of FK506, a cell infected with Ng OE in vehicle only, and a cell infected with Ng OE in the presence of FK506. Downward arrows indicate the timing of STDP induction. Traces show averaged EPSPs indicated with 1 and **2** (scale bars, 2 mV, 50 ms). (**e**) Averaged summary graphs of STDP at 20-ms interval in uninfected control cells (n=7) and Ng OE cells in vehicle (n=6). Each circle represents mean ± SEM. (**f**) Averaged summary graphs of STDP at 20-ms interval in uninfected control cells (n=9) and Ng OE cells in FK506 (n=6). Each circle represents mean ± SEM. (**g**) Collective data of STDP at 20-ms interval in control cells in vehicle (n=7, 105.7 ± 13.3%), control cells in FK506 (n=9, 177.1 ± 16.3%), Ng OE cells in vehicle (n=6, 215.6 ± 24.4%) and Ng OE cells in FK506 (n=6, 204.0 ± 26.9%) cells. EPSP after LTP induction (% baseline) values from individual cells are shown as small filled or open circles. The average values are shown as large filled or open circles with SEM (\*\**p*<0.01, \**p*<0.05, Two-way ANOVA and Tukey’s multiple comparison test).

If Ng OE facilitates the expression of STDP-LTP by inhibiting basal PP2B activity, then FK506 treatment is expected to occlude the facilitative effect of Ng OE on STDP-LTP. We tested this hypothesis by comparing STDP-LTP in Ng OE with and without FK506 treatment. Indeed, the magnitudes of LTP induced by the 20-ms pairing protocol in Ng OE were comparable in the presence (Fig. 7d, f, g) or the absence of FK506 (Fig. 7c, e, g). These results show that both FK506 and Ng OE broaden the temporal window of association for STDP-LTP, and Ng OE facilitates STDP-LTP by suppressing PP2B activity. However, given that FK506 did not alter the decay kinetics of synaptically evoked NMDAR currents in control cells (Fig. 6a), the facilitative effect of Ng OE or FK506 on STDP-LTP may not result from the changes in synaptic NMDAR currents, but from the suppression of PP2B activity on other targets.

Taken together, our results show that Ng controls the induction of spike-timing-dependent LTP at SC-CA1 synapses in the hippocampus, and that decreased Ng levels induce a significant shift in the phosphoproteome signature, enriched by various ASD and schizophrenia targets as well as PSD components. Among the significant targets, we found that hypo-phosphorylation of the NMDAR subunit Grin2A in the C-terminus caused the accelerated decay of NMDAR-mediated currents, highlighting a mechanism for Ng-dependent modulation of synaptic plasticity by regulating PP2B activity.

## Discussion

Translating incoming Ca^2+^ signal into long-lasting changes in excitatory synaptic strength through Ca^2+^/CaM-dependent signaling cascade is essential for information encoding in the brain. Here we show that the levels of Ng, a CaM-binding protein expressed in the postsynaptic compartment of excitatory neurons, control the efficacy of this process through regulating PP2B activity (Supplementary Fig. 4).

In CA1 neurons, the concentrations of Ng and CaM are estimated to be around 20 μM and 10 μM, respectively^49^. Given the tight binding affinity between Ng and CaM^101^, it has been proposed that a majority of CaM is captured by Ng in the postsynaptic compartment in resting neurons^49^. Therefore, endogenous Ng in control cells strictly limits the availability of CaM and its access to Ca^2+^. However, when Ng levels are decreased, more CaM are released from Ng and thus become available for Ca^2+^ binding^102, 103^. Spontaneous neuronal activity causes a fluctuation in the intracellular Ca^2+^ levels within the dendritic spines^104^, and decreased Ng levels in Ng KD promote the formation of a small amount of Ca^2+^/CaM complex. PP2B is preferentially activated over CaMKII when the amount of Ca^2+^/CaM complex is limited because Ca^2+^/CaM complex has a much higher affinity for PP2B compared to CaMKII^45, 98^. Therefore, Ng KD is likely to enhance basal activity of PP2B, causing the accelerated decay of NMDAR-mediated current by dephosphorylation as well as the impairment of LTP (Supplementary Fig. 4a).

On the other hand, increasing Ng levels interfere with the formation of the Ca^2+^/CaM complex by capturing CaM, thus suppressing PP2B activity (Supplementary Fig. 4b). Blockade of PP2B activity by FK506 mimicked and occluded the facilitative effect of Ng OE on STDP-LTP, indicating that FK506 and Ng OE promote LTP through the same biochemical pathway. It is noteworthy that Ng OE did not influence the decay kinetics of synaptic NMDAR-mediated currents (data not shown), similarly to the lack of effect on synaptic NMDAR-mediated currents by FK506 in control cells (Fig. 6a), which implies the lack of noticeable PP2B activity at synapses. Therefore, the facilitative effect of Ng OE or FK506 on STDP-LTP presumably depends on the inhibition of PP2B activity at peri- or extra-synaptic sites. Interestingly, previous studies demonstrated that Ng is localized at a higher concentration in the dendritic spines^29, 105^, and thus the basal PP2B activity is likely to be higher at peri- or extra-synaptic sites compared to the dendritic spines. Importantly, the activation of CaMKII also depends on the formation of Ca^2+^/CaM complexes, but Ng levels may influence the activation of CaMKII to a much lesser degree compared to the case of PP2B. For activation of CaMKII, a much larger amount of Ca^2+^ influx is required due to the weaker binding affinity of Ca^2+^/CaM toward CaMKII. Considering the fact that Ng dissociates from CaM in the presence of high amount of Ca^2+^ (Fig. 2e), when Ca^2+^ influx is large enough to sufficiently activate CaMKII, a majority of CaM becomes available for Ca^2+^ binding regardless of Ng levels.

Our proteomic analyses revealed that Ng KD has profound effect on the phosphoproteome landscape, highlighting a significant bias towards postsynaptic density components. In particular, a decrease in Ng levels leads to changes in phosphorylation patterns of selective ion channels and neurotransmitter receptors, including Grin2A. This finding is consistent with our functional analysis, showing that NMDAR-mediated currents were more transient with Ng KD, which can be rescued by blocking PP2B activity, and that Grin2A mutants disrupting the four phosphorylation sites identified in the phosphoproteome study with Ng KD exhibited fast current decay kinetics compared to the WT. Our data, therefore, provides a mechanistic insight into why decreasing Ng levels in neurons leads to a deficit in STDP-LTP. Heightened postsynaptic PP2B activity dephosphorylates NMDAR subunit Grin2A, thereby accelerates the decay of the NMDAR-mediated synaptic currents, and contributes to the deficit in LTP caused by Ng KD. Interestingly, Ng KD also leads to significant hyper-phosphorylation of certain targets, suggesting that the effect of Ng KD on the phosphoproteome landscape is not a generic consequence of overall increase in phosphatase activities. The functional consequence of the changes in the phosphoproteome beyond LTP will need further exploration.

Previous studies have shown that a change in the phosphorylation status of Grin2A subunits influences the kinetics of synaptic NMDAR currents^88-90^, in addition to other regulatory mechanisms, such as NMDAR Grin2 subunit composition^106-109^ and NMDAR Grin1 (also known as NR1, GluN1) subunit interaction with CaM^85, 86^. However, little is known about the discrete phosphorylation sites in regulating NMDAR properties. Here we demonstrate that Grin2A C-terminal serine residues whose phosphorylation was regulated by Ng levels have a significant impact on the NMDAR current kinetics. Previous work has demonstrated an important role for the C-terminus of Grin2 in regulating NMDAR gating properties^110^. Furthermore, a change in the phosphorylation status of Grin2A subunits influences the kinetics of synaptic NMDAR currents^89, 90^. However, aside from the demonstration of a few putative phosphorylation sites in the proximal C-terminal region of Grin2A involved in regulating NMDAR channel kinetics^88^, identification and functional characterization of C-terminal phosphorylation sites has been lacking. Using quantitative phosphoproteomic approach, we identified phosphorylation sites in the C-terminus of Grin2A. With the high-throughput patch clamping technique, we were able to resolve the roles of C-terminal phosphorylation on NMDAR current kinetics. The similar effect of the SA and SD mutants on NMDAR channel properties suggest that the C-terminal region is stringently regulated by the phosphorylation state to achieve the modulation of channel properties. Given that there is no crystal structure yet available for NMDARs that includes the C-terminus of Grin2A, it is difficult to postulate how phosphorylation at the distal end of the C-terminus impacts the charge distribution and gating. Although many questions still remain, our work clearly demonstrates significant regulatory functions of distal C-terminal phosphorylation of Grin2A subunit on the channel properties of NMDARs.

Taken together, our studies show that Ng levels in the postsynaptic compartment of excitatory synapses dictate the induction of LTP by regulating PP2B activity. We identified one important target, the NMDAR subunit Grin2A downstream from PP2B in the Ng KD condition, contributing to shifts in STDP-LTP caused by changes in Ng levels. The facilitative role of Ng OE in inducing LTP suggests that the rapid increase in Ng translation following neuronal activity will promote the expression of LTP in the population of neurons receiving a similar pattern of excitatory input repeatedly, thus serving as a positive-feedback regulator for LTP and potentially improving the fidelity of memory encoding.

Ng expression is dynamically regulated in neurons at both translational and transcription levels under different behavioral and hormonal states^21, 29, 111-113^. Therefore, the regulation of Ca^2+^-dependent signaling cascade by Ng levels provides a mechanism how the behavioral states of the animals control the efficacy of the information encoding in the brain. Moreover, given the association of Ng with schizophrenia, Jacobsen syndrome and Alzheimer’s Disease^7, 23-25, 114^, and the profound impact of Ng KD on ASD- and schizophrenia-associated gene targets (Fig. 4), our results highlight that the components in the Ca^2+^/CaM-dependent signaling cascade, in particular PP2B and Grin2A, are potential therapeutic targets for cognitive impairment in these diseases.

## Methods

### Animals

7-9 weeks old male C57BL/6 mice (Charles River, USA) were used in electrophysiology experiments. All mice were housed in a pathogen-free, temperature- and humidity-controlled vivarium on a 12-hour light-dark cycle at the Small Animal Facility at the Massachusetts Institute of Technology, and were given ad libitum access to food and water. All procedures related to animal and treatment conformed to the policies of the Committee on Animal Care (CAC) of the Massachusetts Institute of Technology.

### Primary Neuron Cultures

Dissociated cortical neuron culture was prepared from newborn pups of C57BL/6 mice. After dissecting out the cortical areas, the tissues were mildly digested with papain for 20 minutes at 37°C and dissociated with gentle trituration. Following the digestion, cells were plated on poly-D-lysine-coated 12-well plates containing neurobasal media (Invitrogen) supplemented with B27 (Invitrogen). 5-fluoro-2-deoxyuridine was added in culture media at DIV4 to inhibit the growth of glial cells. Neurons were infected with respective lentiviruses at DIV7 and collected at DIV17 for analysis with immunoblot.

### Cell Lines

Constructs for expressing NMDAR subunits were co-transfected with FLP recombinase (pOG44; Thermo) into FlpIn TREx 293 cells (Thermo) and selected for hygromycin resistance (200 μg/mL) to select for and expand small polyclonal pools of single-copy isogenic cell lines. Cells were cultured in customized NEAA-free DMEM/F12 media (Thermo) with 10% FBS, selection antibiotics (200 μg/mL hygromycin and 15 μg/mL blasticidin), and NMDAR inhibitors (1 μg/mL AP5, DCKA, and MK801). For induction, ∼1×10^6^ cells were plated in the absence of selection antibiotics in a 10-cm dish for 3 days and induced with 1 μg/mL doxycycline hyclate (Sigma) for 48 hours in the presence of NMDAR inhibitors and grown to near confluency to improve consistency and reduce variability in recordings (data not shown). Cells were harvested with Accutase (Sigma), suspended in a 1:1 mix of serum free media and pECS-DCF at a concentration of ∼500k/mL, held in a teflon reservoir chilled to 10C, and recorded as soon as possible.

### Cloning of Lentiviral Constructs and Lentivirus Production

The lentiviral transfer vector FUGW and its variant FHUGW were used to create all lentiviral constructs used in this study. The variant FHUGW contains an H1 promoter that drives the expression of an RNAi cassette. In the knockdown experiment, the shRNA targeting Ng mRNA is expressed under the H1 promoter, and eGFP was expressed simultaneously to label infected cells. In studies with Ng overexpression, Ng (Ng-eGFP) or a Ng mutant lacking the CaM-binding IQ motif (NgΔIQ-eGFP) fusion protein was expressed under the ubiquitin promoter.

For the production of these lentiviruses, HEK cells were co-transfected with the lentivirus transfer vectors above, along with the human immunodeficiency virus packing vectors pRSV/REV and pMDLg/pRRE, and the envelope glycoprotein vector VSV-G using FUGENE6 transfection reagent (Roche, Basel, Switzerland). Supernatants of culture media were collected about 60 hours after transfection, and the lentiviral particles were concentrated by centrifuging at 50,000x g. To infect hippocampal CA1 neurons *in vivo*, the concentrated lentiviral particles were infected into the CA1 area of hippocampus bilaterally via stereotaxic surgery. To infect dissociated cortical neuron cultures, 5 μL of concentrated lentivirus was applied into 1 mL of culture media for each well in a 12-well plate.

### Cloning of NMDAR Subunits

Rat Grin1 (NM_017010.2) was cloned downstream of its 31 amino acid signal peptide and EGFP in frame with a P2A peptide fused to rat Grin2A (NM_012573.3). This cassette was then cloned into a modified pFRT-TO single-copy inducible vector (Thermo). Mutations were generated in Grin2A via PCR mutagenesis and subcloned back into the expression construct.

### Validation of Gene Expression by qPCR

∼1×10^6^ cells were collected 48 hours after doxycycline induction, and total RNA was collected using the RNAeasy Plus kit (Qiagen). 1 μg of total RNA was used to generate random hexamer-primed cDNA using the Transcriptor cDNA synthesis kit (Roche). qPCR was done using SYBR Green (Roche) and custom-designed primers, and data was analyzed using the ddCt method, plotting fold change relative to non-induced controls normalized to ACTB. rGrin1-qPCR-F: gtcatcatcctttctgcaagc; rGrin1-qPCR-R: ccagagatctcgcgttcc; rGrin2A-qPCR-F: caaggccagctgctatgg; rGrin2A-qPCR-R: tgccatcccaagtcacatt; ACTB-qPCR-F: CCAACCGCGAGAAGATGA; ACTB-qPCR-R: CCAGAGGCGTACAGGGATAG.

### Immunoblot Analysis

For lentivirus characterization, cortical neuron culture infected with lentivirus was lysed in the Laemmli sample buffer (Bio-Rad) supplemented with 2-mercaptoethanol, PhosSTOP phosphatase inhibitor cocktail (Roche Diagnostics), and the complete mini EDTA-free protease inhibitor (Roche Diagnostics). After boiling at 95°C for 10 minutes, the protein samples were separated on a 10% SDS-PAGE gel. After transferring at 4°C, the membrane (PVDF, Immobilon-FL) was blocked in 5% milk, 0.2% Tween 20, PBS, and incubated with anti-Ng antibody (Millipore, AB5620 or 07-425, 1:1000) and anti-Actin antibody (Sigma, A2228, 1:3000) for one hour at room temperature. The membrane was washed in 5% milk, 0.2% Tween 20, and PBS four times for 10 minutes each. The membrane was then subsequently incubated with the secondary antibodies goat anti-mouse 680 (Licor) or goat anti-rabbit 800 (Licor) conjugated with IR dyes at room temperature for one hour. After washing the membrane, bands were visualized with the Licor Odyssey imaging system.

For the characterization of NMDAR subunits, ∼1×10^6^ cells were collected 48 hours after doxycycline induction and total protein lysate was collected from frozen pellets in lysis buffer (in mM; 1 sodium orthovanadate, 20 sodium phosphate, 5 EDTA, 5 EGTA, 100 sodium chloride, 10 sodium pyrophosphate, 50 sodium fluoride, 1% Triton X100) rocked at 4°C for 1 hour before preclearing. 25 μg of non-boiled protein lysate was run under denaturing conditions on NuPage 3-8% TA precast gels (Thermo), transferred to nitrocellulose membranes (TransBlot, HMW protocol; BioRad), and blotted overnight with primary antibodies against Grin1 (abcam ab109182; 1:1000 in 5% milk), Grin2A (C-term: Millipore 05-901R; N-term: GeneTex GTX103558; 1:1000 in 5% milk), and ACTB (Sigma A5441; 1:50000 in 5% BSA). Western blots were visualized using femto ECL detection (Pierce) with HRP-conjugated secondary antibodies (NA931V 1:10,000, NA9340V 1:2500 (GE)) and documented with a ChemiDoc (BioRad) using exposure times under 10 seconds.

### Phos-Tag SDS-PAGE Analysis

A separating gel with 5% acrylamide was prepared using Tris-HCl solution (pH 8.8) with 25 μM Phos-Tag acrylamide and 50 μM MnCl_2_. The gel was further strengthened by adding 0.6% agarose before the gel polymerization started. A stacking gel with 4% acrylamide was prepared using Tris-HCl solution (pH 6.8) and was strengthened by adding 0.3% agarose. Following the completion of gel running in the Tris-glycine buffer, the gel was soaked in the transfer buffer containing 5 mM EDTA three times for 10 minutes each to remove Mn^2+^ ions from the gel, then washed in a regular transfer buffer without EDTA for 10 minutes. The proteins were transferred to the PVDF membrane (Immobilon-P^SQ^), and after the transfer, the membrane was blocked in 5% BSA, 0.1% Tween 20, and TBS. The membrane was then incubated in a primary antibody, anti-Grin2A (Millipore, 07-632, 1:500), in 5% BSA, 0.1% Tween 20, and TBS overnight at 4°C, then washed in 5% BSA, 0.1% Tween 20, and TBS four times for 10 minutes each. The membrane was subsequently incubated with a secondary antibody goat anti-rabbit 800 (Licor) conjugated with IR dyes at room temperature for one hour. After washing the membrane, bands were visualized with the Licor Odyssey imaging system.

### Pull-down of Ng using CaM Beads

HEK cells were transiently transfected with a plasmid expressing either wildtype Ng (Ng WT) or a mutant form of Ng lacking CaM-binding IQ motif (NgΔIQ) using lipofectamine 2000. Following 20 hours of expression, the cells were cooled down on ice and quickly washed with ice cold PBS. The cells were scraped into cold PBS and pelleted at 3000x g for five minutes at 4°C. After removing the supernatant, the cell pellet was re-suspended and lysed in cold lysis buffer containing 150 mM NaCl, 20 mM Tris (pH 7.5), 1 mM DTT, complete mini EDTA-free protease inhibitor (Roche Diagnostics), 1% Triton X-100 and either 2 mM EGTA or 2 mM Ca^2+^. After spinning down the lysed samples at 10000x g for 10 minutes at 4°C, insoluble precipitates were removed and 10% of the supernatant was saved as an input. The remaining 90% of the supernatant was added to the pre-washed CaM beads (Calmodulin Sepharose 4B, GE Healthcare, 17-0529-01) and incubated overnight with gentle rotation. The CaM beads were pelleted at 2000x g for one minute, and the supernatant was saved as a flow-through portion. The beads were then washed with a lysis buffer three times for 10 minutes each at room temperate, and the bound proteins were eluted from the beads by boiling in 1% SDS for 10 minutes. The amount of Ng in input, flow-through, and CaM bead pull-down fractions were examined using immunoblot analysis.

### Live Cell Imaging

Cells were plated on glass bottom 35-mm tissue culture dishes (MatTek) and induced for 2 days with doxycycline. Cells were maintained in a stage top incubator (okolabs) at 37°C and 5% CO_2_ and imaged with a CSU-X1 spinning disc confocal (Andor) using a Ti-Eclipse microscope and a 60x oil objective (Nikon).

### Stereotaxic Surgery and Preparation of Acute Slice

Stereotaxic surgery was used to inject concentrated lentivirus particles into the CA1 region of the hippocampus in C57BL/6 mice. In this procedure, 7-week-old mice were anesthetized with a ketamine/xylazine cocktail by intraperitoneal injection. After confirming anesthesia, lentivirus particles were injected into the hippocampus based on the antero-posterior and lateral coordinates assigned to the CA1 region. Following the injection, animals were returned to their cages and allowed to recover.

### Electrophysiology

All experiments were performed 5-9 days after stereotaxic injection of lentiviral particles. Acute hippocampal slices (300-μm thick) were prepared based on a published protocol^115^.

STDP experiments were carried out under current-clamp configuration at 30°C in artificial cerebrospinal fluid (ACSF) containing 119 mM NaCl, 2.5 mM KCl, 1 mM NaH_2_PO_4_, 26 mM NaHCO_3_, 11 mM D-glucose, 2.5 mM CaCl_2_, 1.3 mM MgCl_2_, and 100 μM picrotoxin. ACSF was saturated with 95% O_2_ and 5% CO_2_. The patch pipette (4.5-7 MΩ) solution contained 130 mM K-gluconate, 10 mM KCl, 10 mM HEPES, 0.2 mM EGTA, 4 mM MgATP, 0.5 mM NaGTP, and 10 mM sodium phosphocreatine.

The Schaeffer collaterals were stimulated at 0.1 Hz to evoke baseline excitatory postsynaptic potentials (EPSPs) of 3-8 mV. STDP was induced by 100 pairings of presynaptic and postsynaptic stimulations at 5 Hz. Each pairing was consisted of stimulation at Schaeffer collaterals followed by four action potentials given at 100 Hz at various positive time intervals. Each action potential was evoked by injecting a brief depolarizing current pulse (3 ms, 1-2 nA) through the patch pipette. Induction of LTP was monitored over 30 min following the pairing stimulation. All recording data were collected using acute slices prepared from lentivirus-injected animals, either uninfected neurons as a control or infected neurons. Small, hyperpolarizing voltage steps were given at the beginning and end of each recording to monitor input and series resistances under the voltage clamp configuration. In the case of all current clamp experiments, small hyperpolarizing current steps were given for on-line monitoring of input resistance. The cells in which input resistance changed by more than 30% throughout the recording were discarded.

Potential changes in presynaptic release probability were accessed by measuring the PPR. PPR recording was performed under voltage clamp configuration, and targeted neurons were recorded at a holding potential of −70 mV. Excitatory postsynaptic currents (EPSCs) were recorded in response to Schaeffer collateral stimulation. Two consecutive EPSCs were evoked using paired-pulse stimulation with a 50-ms interval, and the recording was repeated 30 times at 0.1 Hz. To measure AMPAR/NMDAR ratio, CA1 neurons were patched under voltage clamp configuration and initially held at −70 mV for 5-10 min to ensure the stability of EPSCs. The cells were subsequently depolarized to +40 mV and EPSCs (mediated by both AMPAR and NMDAR) were monitored for 5-10 min at 0.1 Hz. At that point, D-APV (100 μM) was applied for 10-20 min to isolate AMPAR-mediated EPSCs. A dual component EPSC was obtained by averaging 10-20 consecutive responses immediately before application of D-APV. An average AMPAR EPSC was obtained by averaging 10-20 consecutive responses beginning 7 min after the application of D-APV. An NMDAR EPSC was calculated by subtracting an average AMPAR EPSC from the dual component EPSC. To study NMDAR-mediated current kinetics, EPSCs were recorded in CA1 neurons in a voltage-clamp mode with a −70 mV holding potential. Mg^2+^ was removed from ACSF to unblock NMDAR, and 20 μM CNQX was added in the ACSF to block AMPAR-mediated currents. The decay of NMDAR current was analyzed by fitting the currents to two exponential functions using OriginPro (OriginLab).

In experiments to measure PPR, AMPAR/NMDAR ratio and NMDAR-mediated currents, the patch pipette solution (4.5-7 MΩ) contained 115 mM CsMeSO_3_, 2.8 mM NaCl, 20 mM HEPES, 0.4 mM EGTA, 4 mM MgATP, 0.5 mM NaGTP, 10 mM sodium phosphocreatine, 5 mM TEA-Cl, and 5 mM QX-314. All data were collected using a MultiClamp 700B amplifier (Axon Instruments) digitized at 20 kHz with the analog-to-digital converter ITC-18 computer interface (Heka Instruments). Data were acquired with Igor Pro software (Wavemetrics).

### High-throughput Planar Patch Clamp

All recordings were performed using the SyncroPatch 384PE system (Nanion). External solution was (in mM) 10 HEPES, 80 NaCl, 60 NMDG, 4 KCl, 6 CaCl2, 8 glucose, pH 7.4, 290 mOsm and was supplemented with 30 μM glycine to limit the effects of glycine-dependent desensitization on recordings. Internal solution was (in mM) 20 EGTA, 10 HEPES, 50 CsCl, 10 NaCl, 60 CsF pH 7.2, 285 mOsm. Divalent cation free extracellular solution (pECS DCF) was (in mM) 10 HEPES, 145 NaCl, 4 KCl, 8 glucose, pH 7.4, 300 mOsm. For barium recordings, external solution was the same as above except 6 BaCl_2_ replacing 6 CaCl_2_, and internal solution was (in mM) 10 EGTA, 10 HEPES, 20 CsCl, 90 CsSO4 pH 7.2, 285 mOsm. 10 μM glutamate (sodium salt) was added to external solution for ligand application. Cells were sealed in the whole-cell configuration, held at −60 mV, stimulated with a 5 μL puff of glutamate, and buffer was exchanged 1 sec following ligand application. Recordings were acquired at 5 kHz for 12 seconds post-ligand application. For further details and consideration of SyncroPatch 384PE experimental design, see the reference by Pan et al^116^.

For data analysis, high quality traces were manually selected from DataControl and then analyzed with custom Igor scripts. Briefly, raw traces were read in as waves, Sazitzky filtered with 8 poles at 30 kHz, and trimmed to first 5 seconds of recording which were the most consistent between wells and across biological replicates. Peak currents were extracted before traces were normalized to peak current, fit with a two-exponential for decay kinetics, and averaged. The fast component of the two-exponential was reported as the decay tau, as the slow component was more variable, likely due to the incomplete removal of glutamate after ligand delivery (data not shown). Plots and statistical analysis were done in GraphPad Prism.

### Proteome and Phosphoproteome Study

#### Cellular lysis and enzymatic digestion

Mouse primary neuronal cultures were lysed in 8 M urea with protease and phospho-protease inhibitors, and subsequently digested following a protocol described elsewhere^66^. A small aliquot of cellular lysate was removed from each sample for protein quantification via the Pierce BCA assay kit (Pierce, Rockford, IL). After proteolytic digestion, the samples were quenched with formic acid to a final concentration of 1.0% and subsequently desalted on 30 mg OASIS HLB solid phase columns (Waters, MA, USA).

#### Tryptic peptide labeling with TMT reagent

From each condition (n=6) 460 µg aliquots of the Ng KD dried tryptic peptides were reconstituted in 100 mM HEPES (pH 8.0) to a final concentration of 1.0 mg/mL. The peptides were labeled with Thermo Fisher TMT-6 isobaric mass tag reagent according to manufacturer’s instructions (Thermo Fisher). The peptides were labeled at a 1:8 ratio of peptide to TMT reagent, followed by one hour incubation at room temperature with bench top shaking at 850 rpm. After incubation, a 1.0 µg aliquot of labeled tryptic peptide was removed from each labeled condition, desalted with C18 stage tips^117^, and analyzed by mass spectrometry to ensure that isobaric label incorporation ≥ 95%. An additional 1.0 µg of labeled tryptic peptide was removed from each channel, mixed together, desalted on a C18 stage tip, and analyzed via mass spectrometry to ensure equal relative protein loads. During these quality control steps the labeled peptides were stored, unquenched at −80°C. After validation, each channel was quenched with a 5% hydroxylamine solution to a final sample concentration of 0.3% to quench any unbound isobaric tags. The corresponding 6 channels were mixed together for a total amount of 2.8 mg of labeled tryptic peptides. The labeled peptide mixture was dried down in a speedvac, reconstituted in 500 µl of 3% acetonitrile/0.1% formic acid, and subsequently desalted on tC18 Sep-Pak columns (Waters, MA, USA) in preparation for basic reverse phase fractionation.

#### Basic reverse phase fractionation

The dried peptides were reconstituted in 800 μl of 5 mM ammonium formate (pH 10), and were separated by basic reversed-phase chromatography on an Agilent Zorbax 300 Å 4.6mm x 250mm Extend-C18 column, using an Agilent 1100 Series HPLC instrument (Agilent Technologies). Solvent A (2% acetonitrile, 5 mM ammonium formate, pH 10), and a non-linear increasing concentration of solvent B (90% acetonitrile, 5 mM ammonium formate, pH 10) was used as the mobile phase with a flow rate of 1 ml/min through the column. A non-linear gradient with increasing percentages of solvent B with 4 different slopes was used (0% for 7 min; 0% to 16% in 6 min; 16% to 40% in 60 min; 40% to 44% in 4 min; 44% to 60% in 5 min; 60% for 14 min) and the eluted peptides were collected in a Whatman polypropylene 2 mL 96 well plate. A total of 96 fractionations were collected (∼1 ml/fraction) for a total run time of 96 minutes. The 96 fractions were concatenated into 25 larger fractions, based on the concatenation protocol described elsewhere^118^. From these 25 fractions, 5% of the total volume was removed and used for global proteome analysis. The remaining 95% of each of the fractions were further concatenated down to 13 fractions and phosphopeptide enrichment was performed with these fractions following the IMAC phospho-enrichment protocol described elsewhere^66^.

#### Mass spectrometry analysis

Both the proteome and phosphoproteome were analyzed using a Thermo Fisher Q-Exactive Plus mass spectrometer coupled to a Thermo-Scientific EASY-nLC 1000 liquid chromatograph (Thermo Fisher Scientific). Peptides were separated at a flow rate of 200 nL/min on a self-made capillary column (Picofrit with a 10-μm tip opening and 75 μm diameter, New Objective, PF360-75-10-N-5) packed with 20-cm of C18 1.9 µm silica beads (1.9-μm ReproSil-Pur C18-AQ medium, Dr. Maisch GmbH, r119.aq). Injected peptides were separated at a flow rate of 200 nL/min with a linear 84-min gradient from 100% solvent A (3% acetonitrile, 0.1% formic acid) to 30% solvent B (90% acetonitrile, 0.1% formic acid), followed by a linear 9-min gradient from 30% solvent A to 90% solvent B for a total of 110 minutes. The Q-Exactive plus instrument was operated in the data-dependent mode acquiring higher-energy collisional dissociation tandem mass spectrometry (HCD MS/MS) scans (*R*esolution = 17,500) for TMT-6 on the 12 most abundant ions using an MS1 ion target of 3 × 10^6^ ions and an MS2 target of 5 × 10^4^ ions. The maximum ion time used for the MS/MS scans was 120 ms; the HCD-normalized collision energy was set to 31; the dynamic exclusion time was set to 20 secs, and the peptide-match preferred setting was enabled.

#### Quantitation and identification of peptides and proteins for both proteome and phosphoproteome

All mass spectra were processed using Agilent Spectrum Mill Proteomics Workbench software package pre 6.0 commercial release. For peptide identification, the MS/MS spectra were searched against the UNIPROT mouse database (http://www.uniprot.org/uniprot/?query=proteome:UP000000589) with frequently occurring laboratory contaminants added to the list. The peptides were searched with both fixed and variable modifications. The fixed modifications included N-terminal and lysine modification with the TMT-6 isobaric mass tag and carbamidomethylation turned on. The variable modifications included were N-terminal acetylation, oxidized methionine, and the phosphorylated amino acids serine, threonine, and tyrosine to account for phosphorylation sites. Database matches for the individual spectra were auto-validated by a user-defined threshold for peptides (false discovery rate (FDR) < 1.2%) and an automatic threshold for proteins in a two-step process. In Spectrum Mill, FDRs are calculated at 3 different levels: spectrum, distinct peptide, and distinct protein. Peptide FDRs are calculated in Spectrum Mill using essentially the same pseudo-reversal strategy previously evaluated^119^ and shown to perform the same as concatenation. A false distinct protein identification occurs when all of the relevant peptides, which group together to constitute a distinct protein, have a delta Forward Reverse Score < 0. Spectrum Mill also performs protein grouping using the methods described^120^. Briefly, when a peptide sequence (>8 residues long) is contained in multiple protein entries in the protein database, the proteins are grouped together, and the highest scoring peptide and its accession number are reported. In some cases, when the protein sequences are grouped in this manner, there are distinct peptides which uniquely represent a lower scoring member of the group (isoforms and family members). Each of these instances spawns a subgroup, and multiple subgroups are reported and counted towards the total number of proteins. TMT reporter ion ratios are obtained by calculating the median reporter ion ratio over all distinct peptides assigned to that protein subgroup.

#### Data analysis

The significance of changes in the phosphorylation of individual peptides was evaluated by moderated T-test with a Benjamini-Hochberg correction using the Limma package^121, 122^. Changes in phosphorylation sites on proteins for which total proteome information was available were normalized to the change in total protein quantity as determined by Spectrum Mill. The set of significantly and differentially phosphorylated sites was taken as the union of the sites significant after normalization and non-normalized significant sites for which total protein level quantification was not available. Overlap analysis was performed with this set of proteins against the PSD proteins identified by Bayes et al^73^, with significance evaluated by hypergeometric test. Pathway enrichment analysis was performed on the set of differentially phosphorylated proteins overlapping the PSD using the clusterProfiler R package^76^.

### Statistical Analysis

All bar graphs are presented as the means ± standard error of the mean (SEM). The sample size and statistical methods used in each experiment is provided in the relevant figure legends. All statistical analysis was conducted using GraphPad Prism 7.02 (GraphPad Software Inc.), and significance is shown as *p<0.05, **p<0.01, ***p<0.001.

## Acknowledgements

The authors would like to thank Y. Liu and X. Ren for their excellent technical support, and Drs. E. Scolnick, E. Nedivi, L.-H. Tsai, M. Bear, J. Lisman and M. Wilson for their comments. This work was supported by the Broad Institute Stanley Center Neuropsychiatry Initiative grant, the JPB foundation and the Whitehall foundation (W. X.).

## Author contributions

Conceptualization, H.H. and W.X.; Methodology, H.H., A.A.^5^, H.H.^1^, J.Q. P., S.A.R., R.A. and W.X.; Investigation, H.H, M.J.S, A.A.^5^, A.N.V., N.A.V., S.C.P., and S.Y.W.; Formal Analysis, H.H., L.J.D, F.G., R.A., A.A.^5^, A.A.^6^, and W.X.; Writing – Original Draft, H.H., W.X. and R.A.; Writing – Review & Editing, H.H., W.X., A.A.^6^, and R.A.; Funding Acquisition, W.X.; Supervision, W.X., J.P.Q., S.A.C and R.A.

---

**Supplementary Fig. 1.**
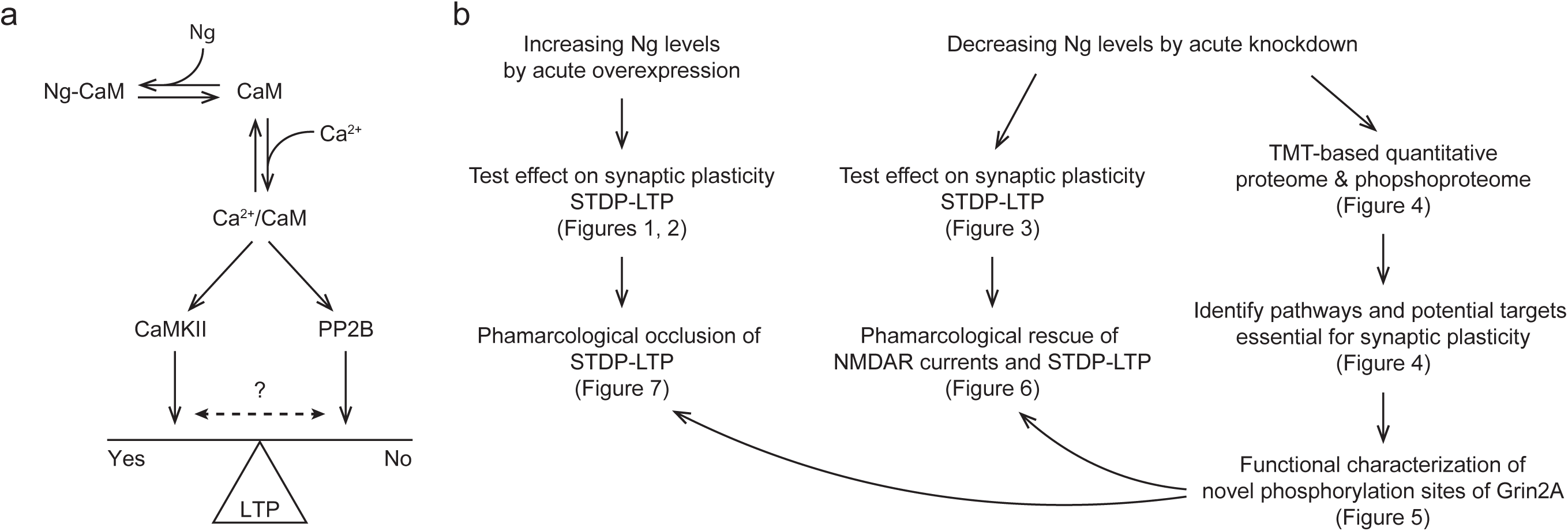
Schematic model of influence of Ng on CaM, and the experimental flow. (**a**) The schematic model of the effect of Ng on relative activation of Ca^2+^/CaM-dependent CaMKII and PP2B that control the expression of LTP. (**b**) The experimental flow. Lentivirus-mediated acute knockdown and overexpression were used to test the effect of Ng levels on synaptic plasticity using STDP-LTP protocol at Schaffer Collateral-CA1 synapses (Figures 1-3). Acute knockdown approach was used for TMT-based quantitative proteomics and phosphoproteomics to analyze how decreasing Ng levels influences the neuronal phosphorylation landscape, and pathway analyses identified potential targets essential for synaptic plasticity, including Grin2A (Figure 4). Interestingly, Ng KD caused hypo-phosphorylation of Grin2A in the C-terminus, and high-throughput planar patch clamp revealed that the phosphorylation sites in the Grin2A C-terminus identified from the phosphoproteome analysis play a critical role in regulating the peak and decay of NMDAR-mediated calcium influx (Figure 5). Lastly, pharmacological approaches were used to validate the potential downstream targets in both the acute knockdown and overexpression conditions (Figures 6, 7).

**Supplementary Fig. 2.**
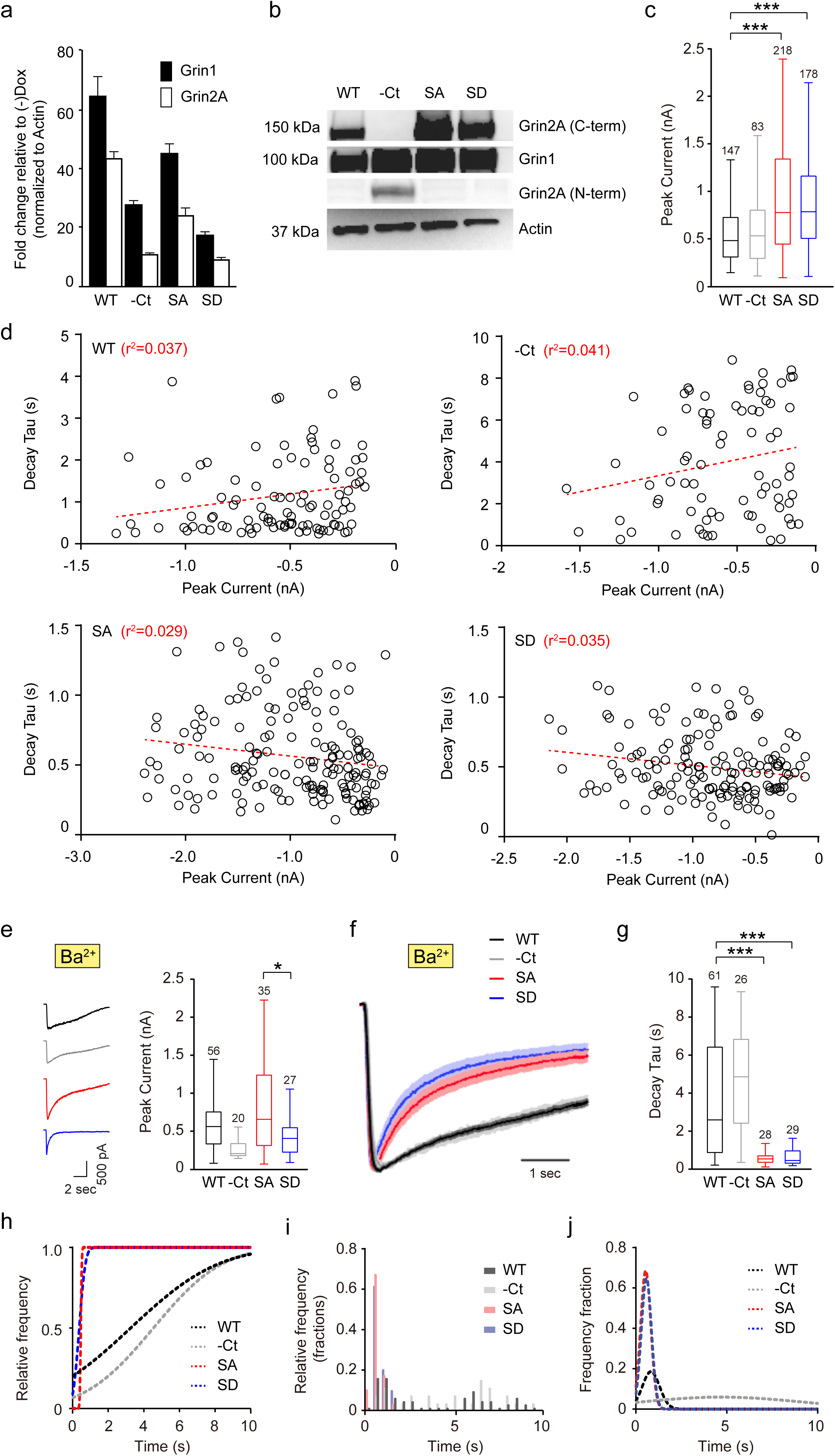
Validation of inducible expression of WT NMDAR and mutants. (**a**) qPCR validation of Grin1 and Grin2A expression two days post-induction with doxycycline. Data are represented as the fold change in mRNA expression compared to non-induced parallel cultures normalized to β-actin. (**b**) Western validation of Grin1 and Grin2A expression two days post induction with doxycycline. The C-terminus deletion mutant was blotted with a polyclonal antibody raised against the N-terminus of Grin2A as the C-terminal epitope recognized by the superior monoclonal antibody is not present in this mutant. (**c**) Box plots of peak current values of NMDAR currents recorded from the cell lines with Grin2A WT, and –Ct, SA and SD mutants. Data were compared via one-way ANOVA and significance was calculated with the Holmes-Sidak multi-comparisons test. \*\*\**p*<0.001. (**d**) Linear regression of the correlation between peak current and decay tau for the recordings from NMDAR WT and mutants show no significant correlation between current size and decay kinetics. Dotted lines represent linear regression fits, and the value of r^2^ for each fit is less than 0.05 demonstrating the poor correlation between current amplitude and decay kinetics. (**e**) Left: Representative recordings of NMDAR-mediated currents recorded in Ba^2+^ containing solution, using planar patch clamp. Right: Box plots of peak current values of NMDAR currents recorded in Ba^2+^ containing solution from the cell lines with Grin2A WT, and –Ct, SA and SD mutants. Data were compared via one-way ANOVA and significance was calculated with the Holmes-Sidak multi-comparisons test. \**p*<0.05. (**f**) Average traces of NMDAR currents recorded in Ba^2+^ containing solution from the cell lines with Grin2A WT and mutants, normalized to peak current highlight differences in decay kinetics in –Ct, SA and SD mutants. Shaded bands represent SEM. (**g**) Box plots of decay Tau values of NMDAR currents recorded recorded in Ba^2+^ containing solution from the cell lines with Grin2A WT and mutants. Data were compared via one-way ANOVA and significance was calculated with the Holmes-Sidak multi-comparisons test. \*\*\**p*<0.001. (**h-j**) Gaussian fits of the cumulative distribution of decay kinetics (h), probability density histograms of decay kinetics (i) and its Gaussian fits (j) from the Ba^2+^ recordings, demonstrate Gaussian distributions for all experimental conditions except for WT and -Ct.

**Supplementary Fig. 3.**
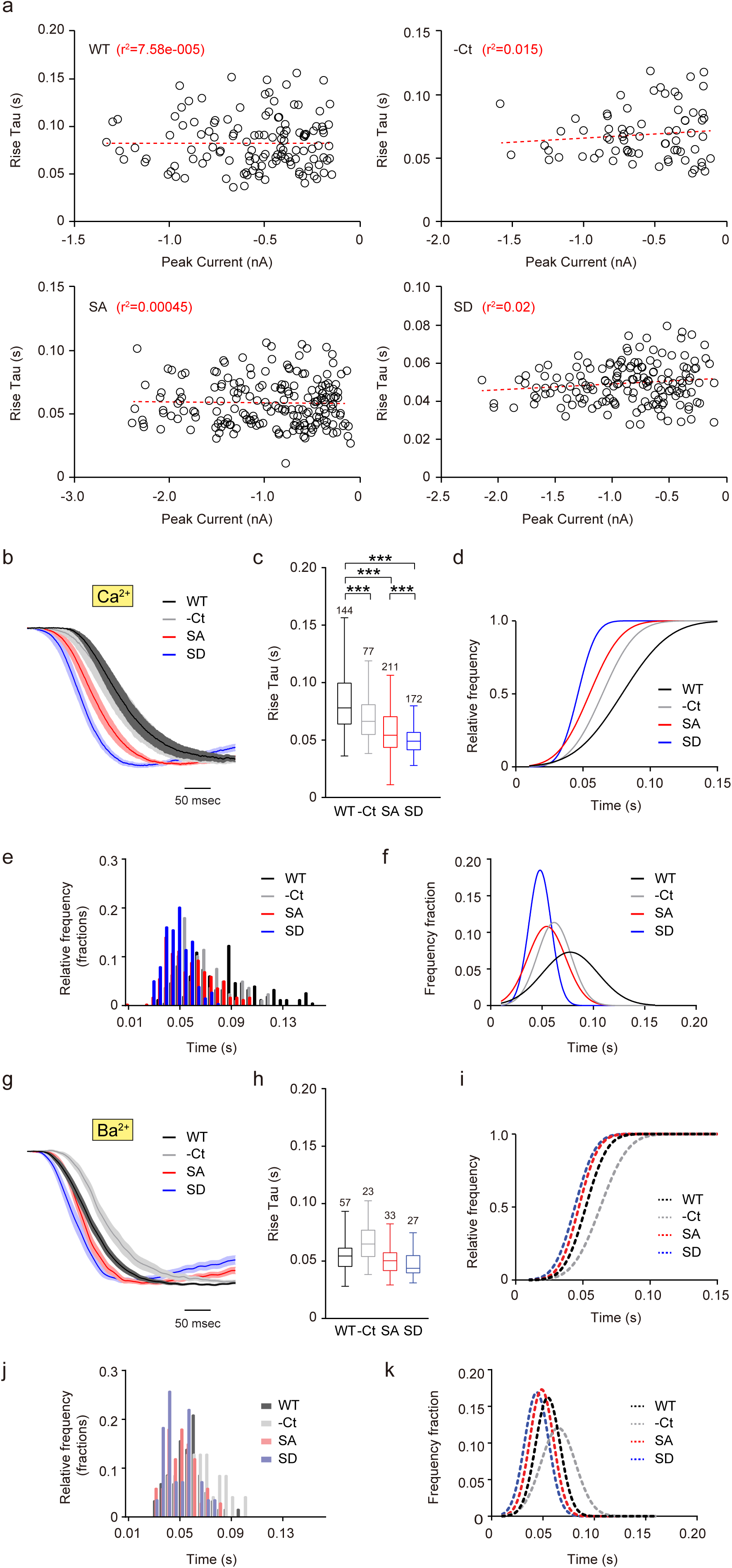
C-terminal phosphorylation of Grin2A regulates the activation kinetics of NMDAR-mediated calcium influx. (**a**) Linear regression of the correlation between peak current and rise tau for the recordings from the cell lines with Grin2A WT and mutants show no significant correlation between current size and rise kinetics. Dotted lines represent linear regression fits, and the value of r^2^ for each fit is less than 0.05 demonstrating the poor correlation between current amplitude and rise kinetics. (**b**) Average traces of NMDAR currents recorded from the cell lines with Grin2A WT and mutants, normalized to peak current highlight the differences in rise kinetics in –Ct, SA and SD mutants. (**c**) Box plots of rise Tau values of NMDAR currents from the cell lines with Grin2A WT and mutants. Data were compared via one-way ANOVA and significance was calculated with the Holmes-Sidak multi-comparisons test. \*\*\**p*<0.001. (**d-f**) Gaussian fits of the cumulative distribution (d), probability density histograms (e) and its Gaussian fits (f) of rise kinetics demonstrates Gaussian distributions for all experimental conditions. (**g**) Average traces of NMDAR currents recorded in Ba^2+^ containing solutions from the cell lines with Grin2A WT and mutants, normalized to peak current highlight the differences in rise kinetics in –Ct, SA and SD mutants. (**h**) Box plots of rise Tau values of NMDAR currents recorded in Ba^2+^ containing solutions from the cell lines with Grin2A WT and mutants. Data were compared via one-way ANOVA and significance was calculated with the Holmes-Sidak multi-comparisons test. (**i-k**) Gaussian fits of the cumulative distribution (i), probability density histograms (j) and its Gaussian fits (k) of rise kinetics in Ba^2+^ condition, demonstrates Gaussian distributions for all experimental conditions.

**Supplementary Fig. 4.**
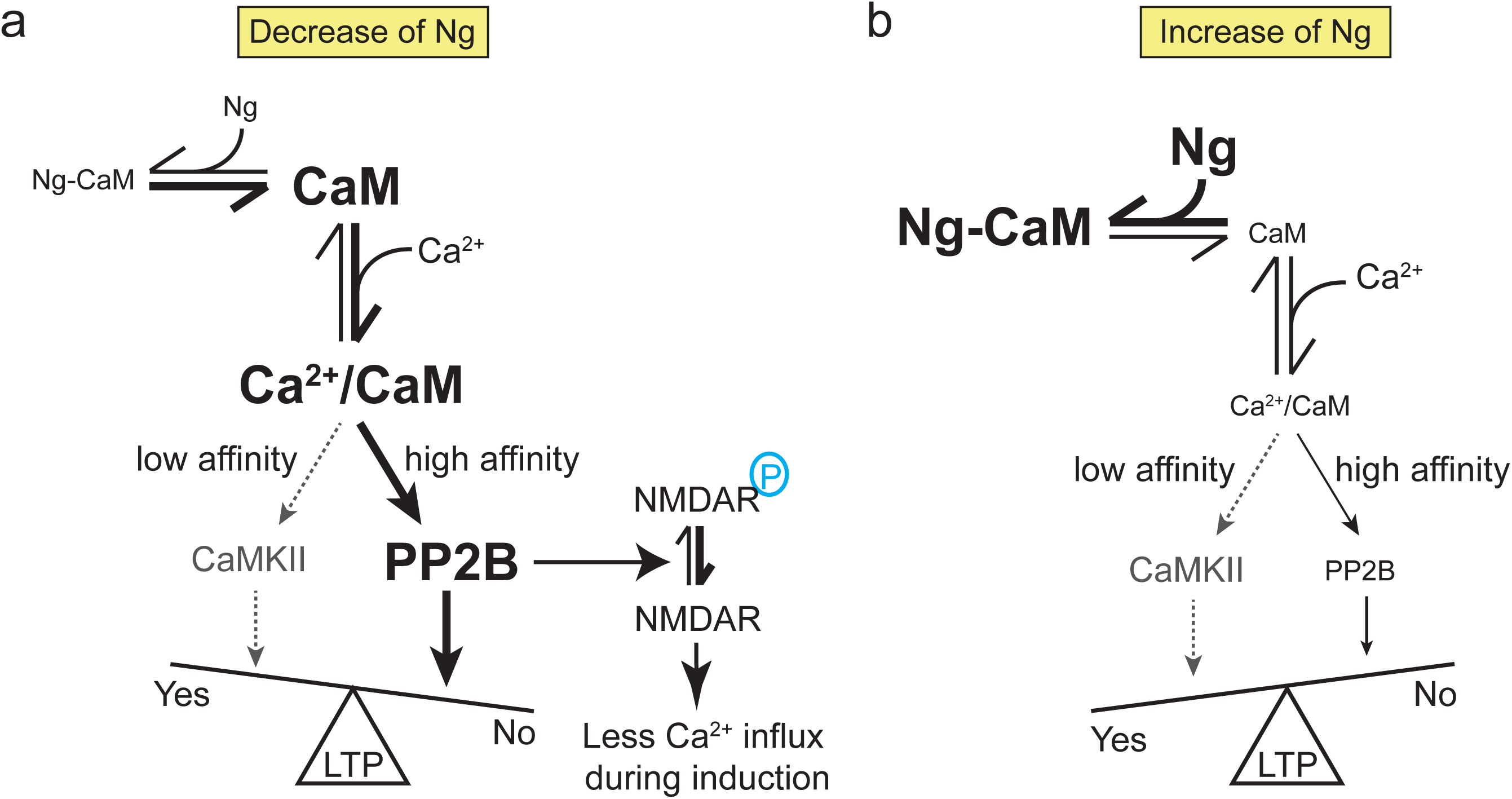
Schematic model demonstrating the effect of Ng levels on LTP. (**a**) Decrease of Ng leads to heightened PP2B activity in neurons, which dephosphorylates the NMDAR subunit Grin2A, and accelerates the decay of synaptic NMDAR currents, thus decreasing Ca^2+^ influx through NMDARs. The changes in this cascade leads to deficit in LTP. (**b**) Increase of Ng decreases basal PP2B activity and lowers the threshold of LTP.

**Supplementary Table 1.**
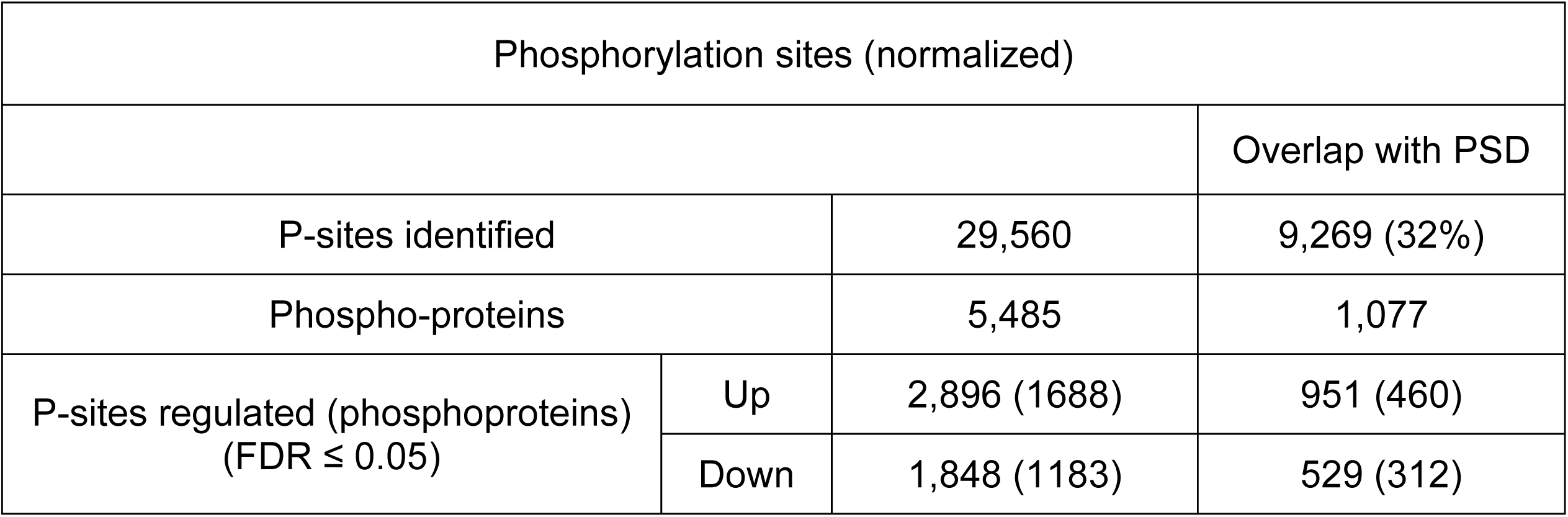
Phosphorylation sites (P-sites) coverage with Ng knockdown.

| Phosphorylation sites (normalized) |  |  |  |
| --- | --- | --- | --- |
|  |  |  | Overlap with PSD |
| P-sites identified |  | 29,560 | 9,269 (32%) |
| Phospho-proteins |  | 5,485 | 1,077 |
| P-sites regulated (phosphoproteins)<br>(FDR ≤ 0.05) | Up | 2,896 (1688) | 951 (460) |
|  | Down | 1,848 (1183) | 529 (312) |

**Supplementary Table 2.**
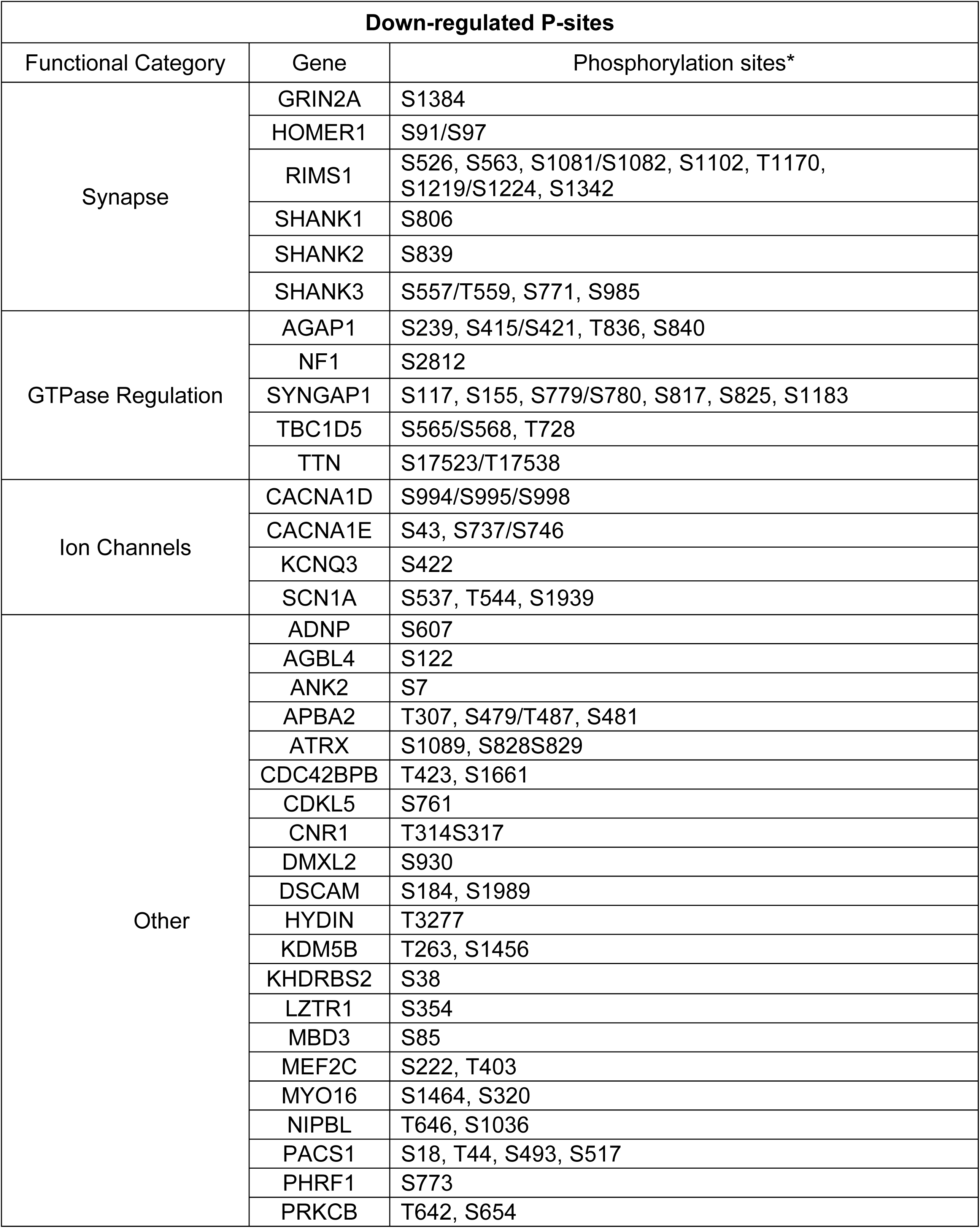
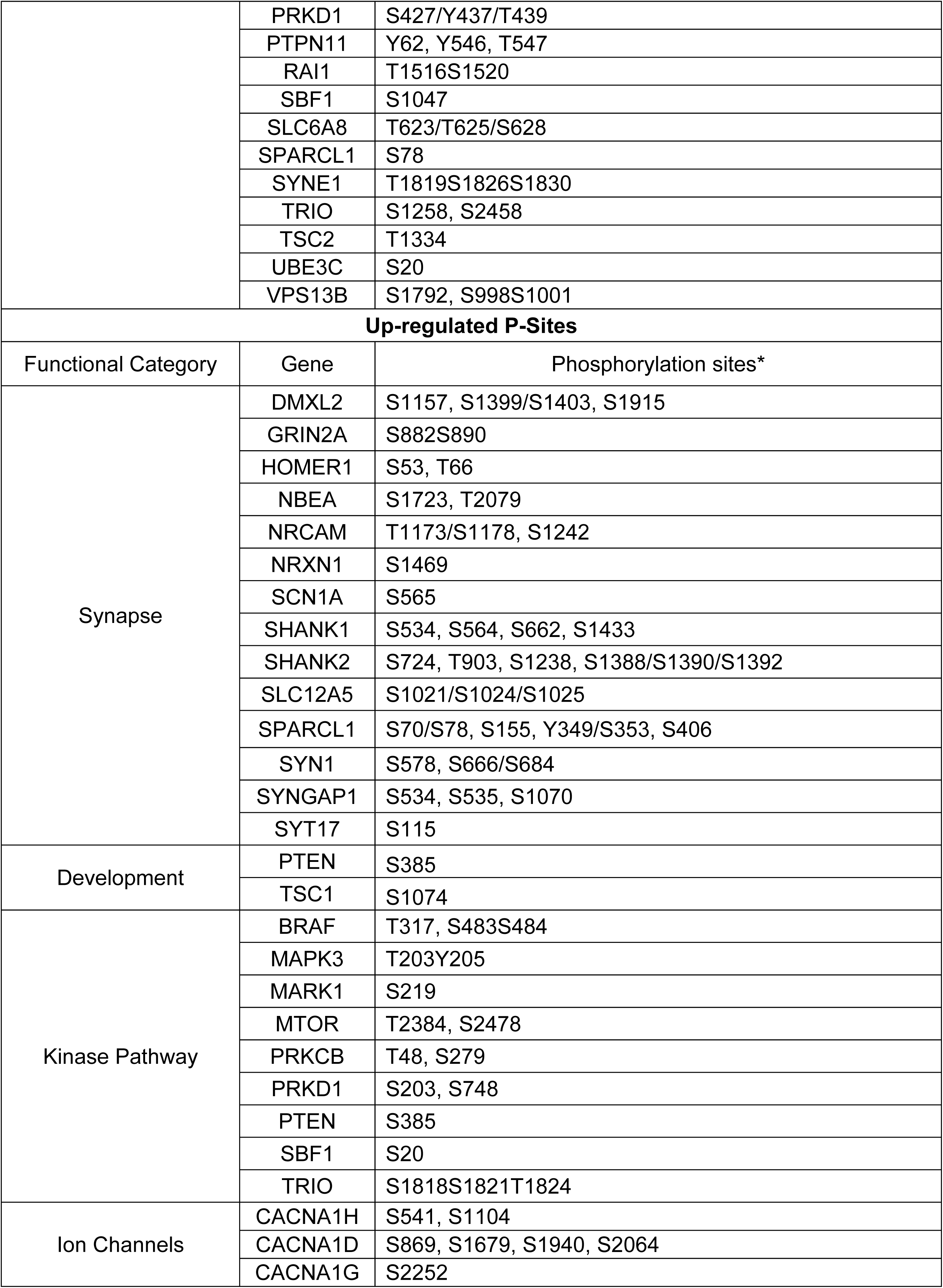
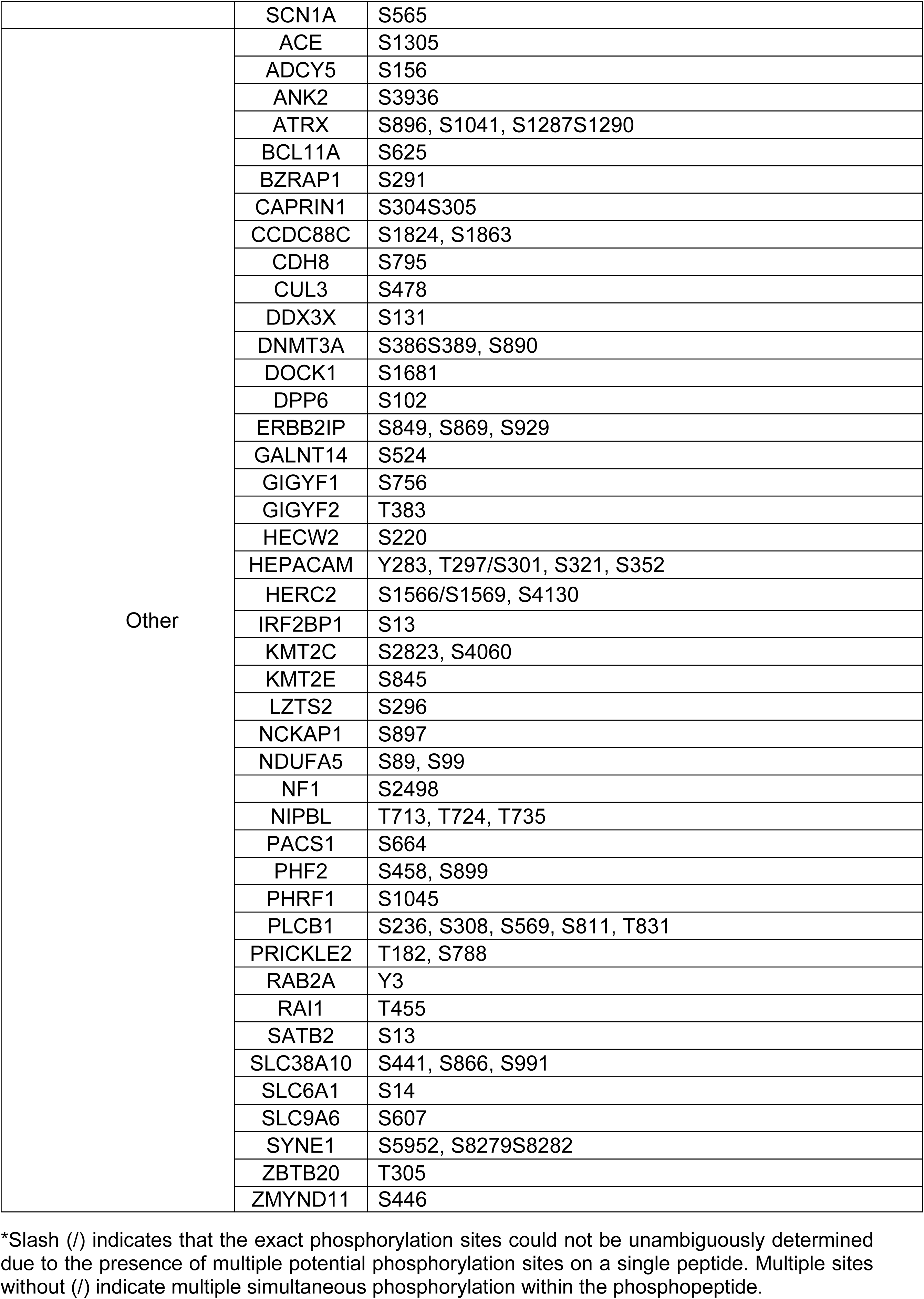
Significantly regulated P-sites in ASD targets affected by Ng knockdown.

**Supplementary Table 3.** Significantly regulated P-sites in SCZ targets affected by Ng knockdown.

|  | Gene | Phosphorylation sites* |
| --- | --- | --- |
| Down-regulated P-sites | ALDOA | S414 |
|  | HCN1 | S99 |
|  | GRIN2A | S1384 |
|  | SMG6 | S239 |
|  | SNAP91 | T788 |
|  | PRKD1 | S427/Y438/T439 |
|  | RANBP10 | S365/S367, S369 |
|  | TBC1D5 | T728, S565/S568 |
|  | CNKSR2 | S109, S488, S928, S936 |
|  | HIRIP3 | T391 |
|  | KDM4A | T361 |
|  | BAG5 | S298 |
|  | ZDHHC5 | S415, S577 |
|  | TMX2 | S187 |
|  | TCF20 | S567, S1395 |
|  | R3HDM2 | S347, S383/T389 |
|  | PLCL1 | T94/S96 |
|  | DOC2A | S226 |
| Up-regulated P-sites | ALDOA | S90 |
|  | ATP2A2 | S186, S378, T441, T537, S553 |
|  | GRIN2A | S882/S890 |
|  | INA | S274, S441 |
|  | VPS45 | T540 |
|  | PLCL1 | S1080 |
|  | SNAP91 | S248, S296/S300/S303 |
|  | PRKD1 | S203, S718 |
|  | MAPK3 | T203/Y205 |
|  | GIGYF2 | T383 |
|  | CNKSR2 | S12, S325/T332/S338, S430 |
|  | SATB2 | S13 |
|  | TMX2 | S274 |
|  | MPP6 | S197 |
|  | CUL3 | S478 |
|  | RFTN2 | S429 |
|  | PSMA4 | T185 |

\*Slash (/) indicates that the exact phosphorylation sites could not be unambiguously determined due to the presence of multiple potential phosphorylation sites on a single peptide. Multiple sites without (/) indicate multiple simultaneous phosphorylation within the phosphopeptide.

